# Local desensitization to dopamine devalues recurring behavior

**DOI:** 10.1101/2024.02.20.581276

**Authors:** Lauren E. Miner, Aditya K. Gautham, Michael A. Crickmore

## Abstract

Goal achievement adjusts the relative importance of future behaviors. We use Drosophila to study this form of motivational control, finding that prior matings make males increasingly likely to abandon future copulations when challenged. Repetition-induced devaluation results from a reduction in dopamine reception by the Copulation Decision Neurons (CDNs), which mediate the decision to end matings. Dopamine signaling to the CDNs sustains matings in real time, but also triggers a lasting, β-arrestin-dependent desensitization of the D2R on the CDNs, leaving subsequent matings susceptible to disruption. When D2R desensitization is experimentally prevented, the male treats each mating as if it were his first. These findings provide a generalizable mechanism of motivational control and reveal a natural function for the long-studied susceptibility of the D2R to drug-induced inactivation.

## Introduction

Each moment an animal spends in a behavior is the outcome of a decision that weighs current and past experiences. Dopamine marks lived experience by creating internal representations of rewards and failures^1– 4^, and by sculpting motivational states that reflect the urgency of the behaviors under consideration^5–9^. How dopamine signaling changes with experience to guide future behavior is therefore a central question in understanding behavioral control.

Many goal-directed behaviors can be separated into appetitive and consummatory phases^10–13^. During appetitive behaviors, animals invest time and energy toward a possible reward and, when successful, enable a transition into the consummatory phase. Though they rise and fall in tandem, appetitive and consummatory behaviors often require a different set of coordinated actions (e.g., foraging vs. feeding), and investigations into mammalian feeding^10,14–16^ and Drosophila mating behaviors^6,17^ have shown their motivations to be functionally separable.

Male flies use dopamine to implement both appetitive and consummatory aspects of mating drive, with anatomically distinct dopaminergic populations that set the tendency to court females^6,18,19^ and, once copulation has been achieved, then set the persistence to continue mating through challenges^17^. While high dopamine levels in the brain can sustain the motivation to court for weeks^6^ (our observations), we find that the dopamine produced in the ventral nervous system that supports males’ perseverance during copulation desensitizes its target neurons, making subsequent matings more likely to be abandoned when challenged.

When left unperturbed, Drosophila copulation lasts ∼23 minutes, with no obvious indication of internal state changes as the mating progresses^17^. But subjecting the male to threats at varying times into mating reveals an increasing susceptibility to abandonment later into mating^17^. Local dopamine signaling counters challenges to mating by dampening the output of the Copulation Decision Neurons (CDNs), whose activity is required to end matings^17,20^. We show that the mating-protective influence of dopamine is conveyed directly to the CDNs through the D2 receptor, an effect that both protects the current mating and desensitizes the receptor for future matings. Males lacking the D2 receptor—or who have recently mated—are less responsive to the motivating dopamine, and, consequently, show an increased likelihood to end matings when challenged. Inversely, flies that were prevented from releasing dopamine during earlier matings, or who lack β-arrestin in the CDNs, endure challenges to repeated matings with the same resiliency as their first.

The D2 receptor is notoriously prone to desensitization in response to pharmaceutical^21^, drug-induced^22–30^, or experimental^31,32^ increases in dopamine. Our results provide a natural function for this long-studied effect, one that is likely to generalize across animals and behaviors. In light of these findings, the broad devaluation of previously rewarding behaviors in addiction may be seen as a widespread corruption of local mechanisms for deprioritizing repeated behaviors.

## Results

### Future copulations are devalued by prior matings

For the first five minutes of a ∼23-minute mating, male flies demonstrate an apparently insurmountable motivation to persist in the face of even lethal challenges^17,33^. As the mating progresses their resiliency declines, such that by 15 minutes even relatively modest environmental changes can truncate most matings^17,20^. If supplied with an abundance of females and left to mate *ad libitum*, males will mate multiple times, depleting their reserves of reproductive fluid after 3-5 matings and rendering future matings unproductive^6,34^. Though courtship decreases after each mating (**Supplementary Figure 1A**), even completely satiated males will still occasionally court and mate^18^, allowing us to ask whether these unproductive matings are guided by the same motivational dynamics as productive ones^17,20^. We found that, after spending 2.5 hours mating *ad libitum* with females in a satiety assay (1 male with ∼15 virgin females^6,34^; see **Methods**), males become much more likely to terminate subsequent matings when challenged (**Figure 1A, B, Supplementary Figure 1B**). This manifestation of satiety lasts for at least a day (**Figure 1B**), holds across multiple threat modalities (**Figure 1C**), and, in agreement with previous reports^6,35^, does not alter the duration of unchallenged matings (**Supplementary Figure 1C**).

**Figure 1:**
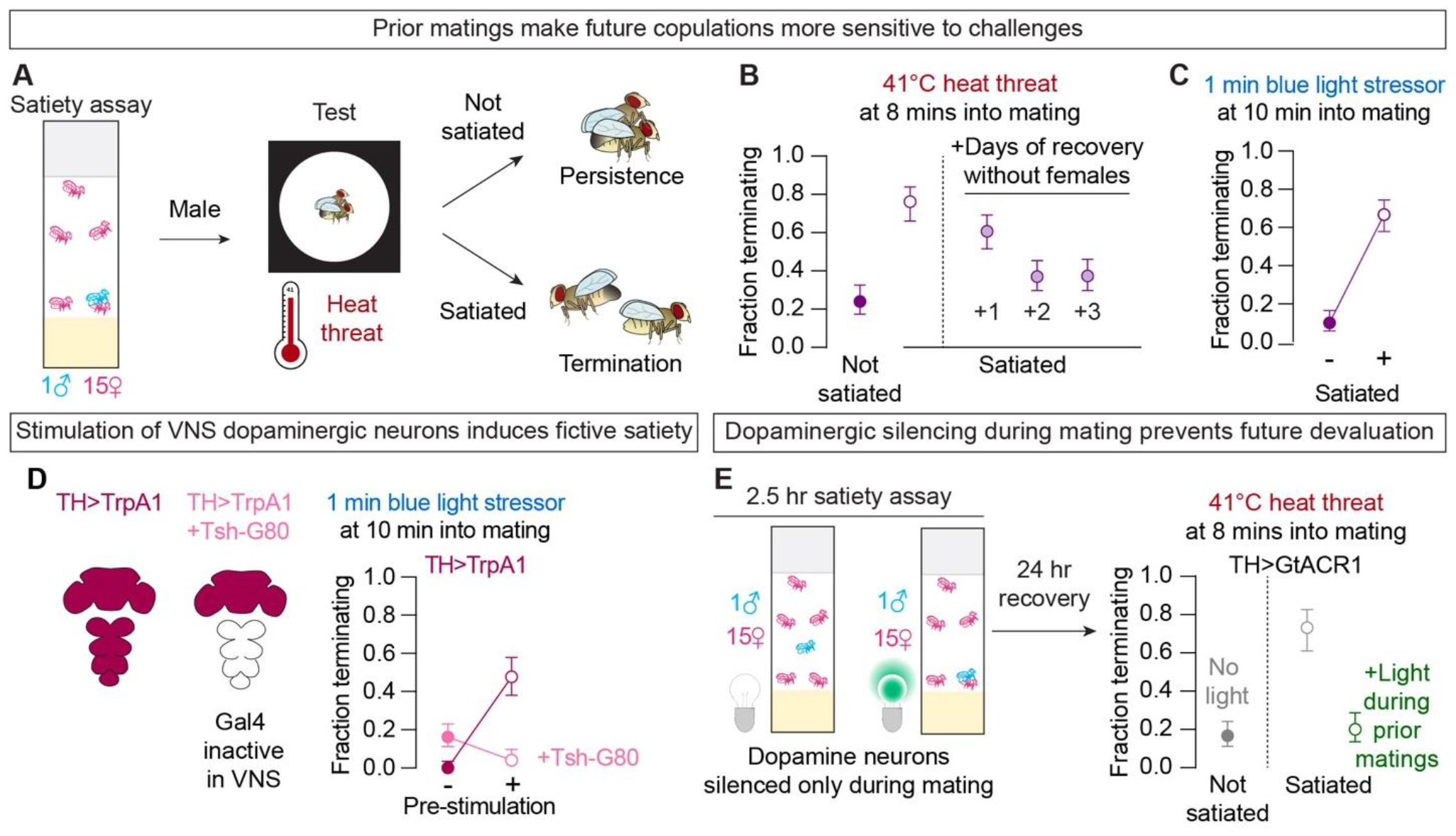
Dopamine released during copulation decreases the resiliency of future matings. **A**. In the satiety assay, individual male flies are provided *ad libitum* access to ∼15 virgin females in a standard food vial^6,34^. After the assay, the males are removed and their motivation to sustain a subsequent mating is tested with a minute-long heat threat. Unless otherwise noted (e.g., **Figure 1C**) we use 2.5 hr satiety assays for all behavioral experiments (see **Methods**). We assess devaluation using threat severities and timepoints with low, but non-zero termination rates in naïve males. **B**. Prior matings increase the probability that a wildtype male will terminate a subsequent mating in response to a 1-minute heat threat delivered 8 minutes after the mating began, an effect that lasts for at least 24 hours (n=21-35). Here and below, mean ± 67% credible intervals are shown for proportions, selected to approximate SEM. Statistical tests and p values are reported for all experiments in **Table S1**. **C**. Recent matings increase the probability that a control (UAS-TrpA1/+) male will terminate a subsequent mating in response to bright blue light (n=30 each). We delivered the bright blue light at 10 min into mating to generate termination probabilities similar to flies receiving a 41°C heat threat at 8 minutes into mating. Flies were satiated for 4.5 hrs to match the length of the pre-stimulation treatment in **Figure 1D** and **Supplementary Figure 2**. **D**. Transient stimulation of dopamine neurons (labeled by TH-Gal4) for 4.5 hrs prior to mating using the warmth-activated cation channel TrpA1 increases the probability that a male will terminate a mating in response to bright blue light (n=23-30). This effect is prevented by the addition of a Tsh-G80 transgene that blocks Gal4 activity in the VNS. **E**. Silencing dopaminergic neurons during prior matings using the green light-gated chloride channel GtACR1 prevents the increased sensitivity to threats during subsequent matings (n=15-30; see **Methods**). Though the number of matings in the satiety assay was unchanged by TH>GtACR1 silencing (**Supplementary Figure 3D**), subsequent courtship was decreased in the 1 male 1 female copulation chambers, so we performed the heat threat experiments after a day of recovery. Detailed genotypes for all experiments are reported in **Table S7**.

As with courtship satiety^6^, it is mating that triggers the devaluation of copulation: no effect is seen after housing the male with unreceptive females (**Supplementary Figure 1D**) or artificial depletion of the male’s reproductive stores (**Supplementary Figure 1E-G**). Blocking sperm transfer during prior matings does not prevent the consequent decrease in threat resiliency (**Supplementary Figure 1H, I**), demonstrating that, as with courtship, the value of copulation is not a function of the true fertility state of the male but results from the lingering effects of previous matings in the nervous system. These results establish a new system for studying the effects of prior experience on motivational dynamics using the accessible and increasingly well-understood sexually dimorphic circuitry of Drosophila.

### Dopamine released during copulation devalues subsequent matings

Much of the circuitry that regulates courtship has been identified^6,18,19,36–38^. While testing courtship circuit elements for potential roles in copulation devaluation (**Supplementary Figure 2**), we made a key observation: after the dopaminergic neurons of non-mating males were transiently stimulated using the warmth-activated cation channel TrpA1, subsequent matings became much more susceptible to threats (**Figure 1D, Supplementary Figure 3A-C**). This effect is essentially the opposite of the mating-protective effect observed when dopaminergic neurons are stimulated *during* (as opposed to *before*) mating^17^ (**Supplementary Figure 3D, E**). Like the motivating effect, this post-stimulation de-motivating effect was blocked by a Tsh-Gal80 transgene that allows activity of Gal4 in the brain but prevents activity in the ventral nervous system ^17^ (VNS; **Figure 1D**), suggesting that the same VNS dopamine neurons are responsible for both the real-time motivating, and the long-term de-motivating effects. The Tsh-Gal80 result also shows that the dopaminergic activity in the brain, which is unaffected by this transgene and governs courtship decisions, has no bearing on motivational dynamics during copulation, reinforcing the idea of anatomical separation of appetitive and consummatory regulation in mating behavior.

The paradoxical effects of dopaminergic stimulation led to the central hypothesis of this work: that dopamine released during the current mating induces a long-lasting devaluation of future matings. To prevent dopamine release during matings we used the green-light gated chloride channel GtACR1^39,40^ to transiently hyperpolarize dopamine neurons. When this silencing was time-locked to copulation, it prevented the devaluation of subsequent matings without changing their duration, whereas the same silencing frequency had no effect when temporally uncoupled from copulation (**Figure 1E, Supplementary Figure 3F-H**). In other words, were it not for the dopamine released during earlier matings, every mating would be treated as the first.

### Dopamine provides a motivating signal to the Copulation Decision Neurons

To understand how previously motivating dopamine devalues future matings, we first asked how and where the signal is received. The original discovery that dopamine is used to set the persistence of the male to continue copulating through challenges^17^ was made before tools were available for inhibitory optogenetics in Drosophila. So we first used GtACR1 to confirm that dopaminergic activity is required to protect copulation from challenges in real time (**Figure 2A**). Next we looked to identify which of the four known dopamine receptors in Drosophila could mediate this protective effect. Of these, mutation of D2R, the sole D2-like receptor in Drosophila, most strongly increased the termination frequency when matings were challenged with heat threats (**Supplementary Figure 4A, B**). This finding gave us the opportunity to identify the downstream targets of dopamine signaling that set the value of continuing to mate when presented with challenges.

**Figure 2:**
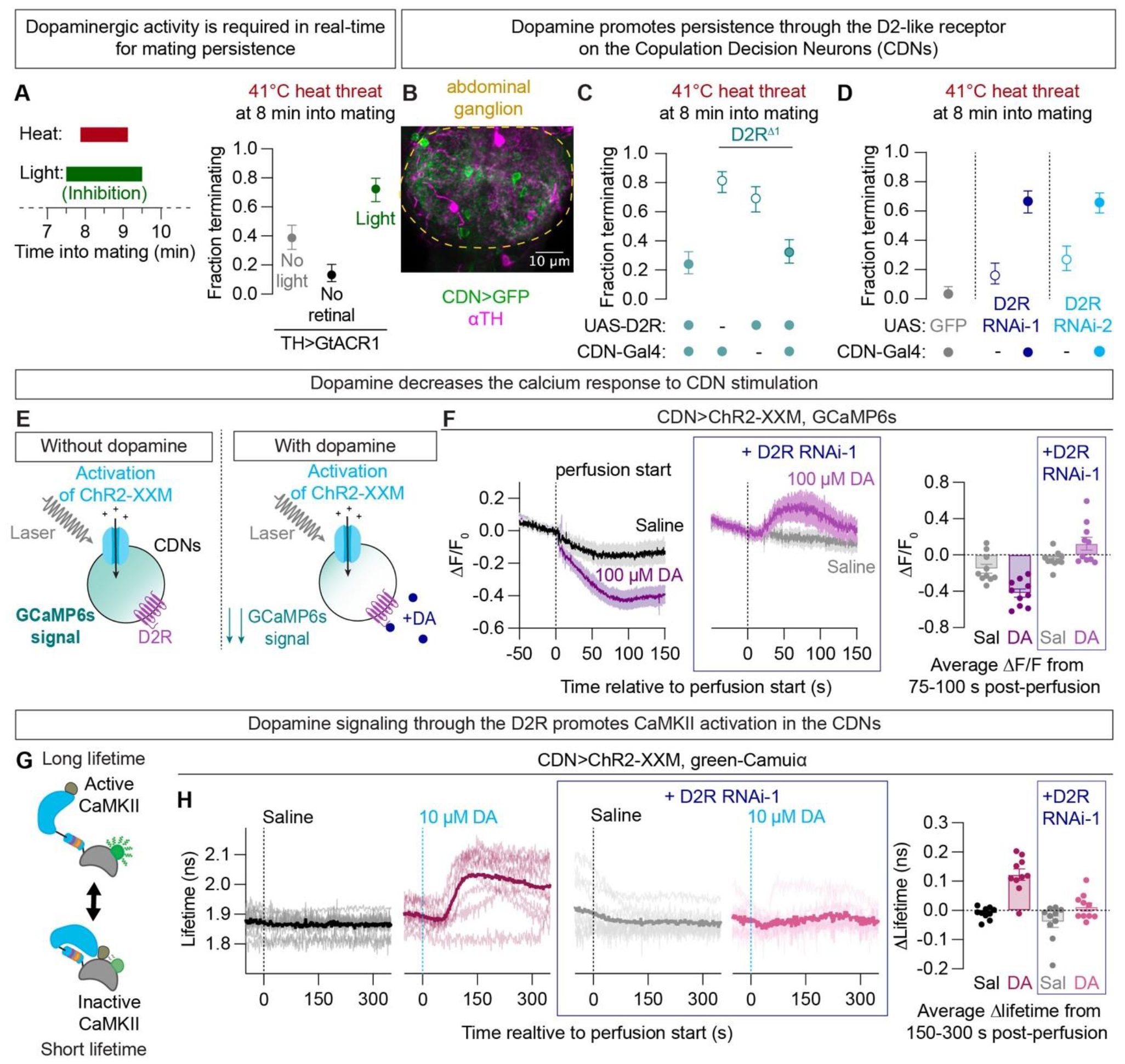
Dopamine protects mating by signaling through the D2 receptor to the Copulation Decision Neurons. **A**.Electrically silencing dopaminergic neurons using GtACR1 from 30s before until 30s after a heat threat at 8 minutes into mating reduces the probability of persisting through the threat, compared to animals that were not fed the obligate chromophore retinal, or trials in which the light was never switched on (n=29-31). We believe the no light control is more sensitive to threats due to partial silencing in low lighting conditions. **B**.Staining for tyrosine hydroxylase in the abdominal ganglion shows dopaminergic neurons closely intermingled with the CDNs. **C**.D2R mutant animals have decreased mating persistence when challenged by threats. Restoring expression of the D2R in the CDNs of D2R mutant animals rescues the phenotype (n=26-31). **D**.RNAi-mediated knockdown of the D2R in the CDNs increases the probability that a male will terminate a mating in response to a heat threat (n=24-44). **E**.Schematic of our explant preparation in which the CDNs are stimulated by ChR2-XXM activated by our laser at ∼10µW, while GCaMP6s is used to monitor calcium dynamics. **F**.Left: Bath application of 100 µM dopamine (DA) decreases the calcium levels caused by optogenetic activation of the CDNs (n=10 each). Trace averages are shown as the mean ± SEM for GCaMP experiments. See Supplementary Figures for whole traces of all GCaMP experiments. Middle: Knocking down the D2R in the CDNs prevents the suppressive effects of dopamine (n=10 each), perhaps even allowing dopamine to drive a net influx of calcium—presumably through D1-like receptors. Right: summary of the GCaMP data with each dot representing the average change in fluorescence between 75 and 100 s post-perfusion in one animal. Bars report the population average as mean ± SEM for all GCaMP experiments. **G**.Green-Camuiα reports CaMKII activity. A conformational change that opens the kinase disrupts FRET, causing an increase in fluorescence lifetime. **H**.Left: In the absence of externally supplied dopamine, optogenetic stimulation of the CDNs does not change CaMKII activity (n=10 each). Right: Bath application of 10 µM dopamine allows the Chr2-XXM stimulation to activate CaMKII (n=10 each). Individual traces shown with average traces bolded. Middle: knocking down the D2R prevents the effects of dopamine on CaMKII activity (n=10 each). Right: summary data with each dot representing the average change from baseline lifetime between 150 and 300 s post-perfusion in one animal. Bars report the population average as mean ± SEM for all Green-Camuiα experiments.

A strong candidate for receiving the signal through the D2R is a set of eight GABAergic Copulation Decision Neurons (CDNs, labeled by NP2719-Gal4), that reside in the abdominal ganglion, in close proximity to dopaminergic neurons (**Figure 2B**)^17^. As expected from neurons that mediate the decision to end matings^17,20^, electrical silencing of the CDNs suppressed the increased termination caused by electrical silencing of dopamine neurons (**Supplementary Figure 4C**). We found that rescuing D2R expression only in the CDNs of D2R mutant animals restored normal threat resiliency (**Figure 2C, Supplementary Figure 4D**), without causing an increase in baseline (i.e., first mating) persistence (**Supplementary Figure 4E**). Inversely, knockdown of D2R in the CDNs using two independent RNAi lines increased the responsiveness to threats (**Figure 2D, Supplementary Figure 4F**). We confirmed the expression of D2R in the CDNs using a recently published single cell sequencing data set (see **Methods**)^41^. Together, these data support a model in which dopamine motivates mating by acting directly on the CDNs through the D2R.

The opposing real-time effects of the CDNs and dopaminergic neurons suggest that dopamine signaling through the D2R may dampen the responsiveness of the CDNs. We used an *ex vivo* preparation in which we stimulate the CDNs while monitoring their calcium dynamics in response to bath application of dopamine. Because the CDNs have low basal activity in this preparation, we expressed the channelrhodopsin variant ChR2-XXM^42^, which we previously noticed was activated by high (∼10 mW) power of our near infrared laser— allowing simultaneous stimulation and imaging (Gautham et al. in revision; **Figure 2E, Supplementary Figure 5A**). Once GCaMP6s levels stabilized, we perfused dopamine into the bath and saw a decrease in calcium levels (**Figure 2E, F; Supplementary Figure 5B, C**), in accordance with the opposing roles of dopaminergic and CDN activity on behavior. As expected, mutating the D2R or knocking it down in the CDNs completely prevented the reduction in calcium levels, even at relatively high dopamine concentrations (100 µM) (**Figure 2F, Supplementary Figure 5D-F**).

In addition to the impact on calcium levels, we recently found a CaMKII-based mechanism through which dopamine impacts CDN function (Gautham et al. in revision). CaMKII activity protects matings from within the CDNs by limiting the duration over which stimulatory inputs can accumulate towards the decision to end the mating (Gautham et al. in revision). Here, the CaMKII response to dopamine provides a molecular readout of dopamine signaling that is compatible with 2-photon fluorescence lifetime imaging (2p-FLIM), a sensitive and robust technique that computes an estimate of fluorescence lifetime by tracking photon arrival times at the photomultiplier tube (instead of simply reporting bulk photon counts)^43^. Importantly, this method provides an absolute measure of CaMKII activity and so does not require normalization to baseline, facilitating comparisons across animals. We used the FRET sensor green-Camuiα^33,44^ to monitor CaMKII activity (**Figure 2G**) under similar conditions as described above and reproduced the finding that the coincidence of gentle ChR2-XXM activation and bath application of dopamine increases CaMKII activity in the CDNs (Gautham et al. in revision; **Figure 2H, Supplementary Figure 5G**, see **Methods** for details). Mutating the D2R or knocking it down in the CDNs reduced the ability of 10 µM dopamine to activate CaMKII (**Figure 2I, Supplementary Figure 5H, I**), indicating a requirement for the D2R to implement both known physiological consequences of the motivating dopamine signal on the CDNs.

### Matings cause a lasting insensitivity to dopamine

We next examined the changes that prior matings impose on the CDNs. We first confirmed that electrical silencing of the CDNs prevented satiated males from ending matings in response to a heat threat at 8 minutes (**Supplementary Figure 6A**), consistent with their role as arbiters of the decision-making process. Optogenetic stimulation of the CDNs causes mating termination with probabilities that depend on stimulation duration and intensity, as well as the time into mating when the stimulus is delivered^17,20^ (Gautham et al, in revision). While basal activity of the CDNs was unchanged (**Supplementary Figure 6B**), we found that prior matings increased the ability of optogenetic CDN stimulation to terminate matings over a range of light intensities, with a magnitude comparable to heat threats (**Supplementary Figure 6C, D**). We saw a similar potentiation of the response to CDN stimulation when the D2R was reduced in the CDNs (**Supplementary Figure 6E**). These data argue that the increased sensitivity to challenges in satiated males can be localized to and understood as an increase in termination-promoting output in response to the same termination-promoting CDN inputs.

One explanation of these results might be that less dopamine is released on repeated matings. To test this idea, we artificially activated the dopaminergic neurons of satiated flies. Dopaminergic stimulation dramatically extends the mating duration of naïve males^17^, but this effect was largely suppressed in satiated males (**Figure 3A, Supplementary Figure 7A**). This result argues for a satiety-induced change in the dopamine-receiving, as opposed to dopamine-releasing, neurons. Consistent with this idea, calcium imaging found no satiety-induced differences in basal or evoked activity of the dopaminergic projections in the abdominal ganglion (**Supplementary Figure 7B, C**). While these experiments are not conclusive, they suggest that decreased CDNs responsiveness to dopamine—not decreased dopaminergic activity—is responsible for the effects of satiety on decision making during mating.

**Figure 3:**
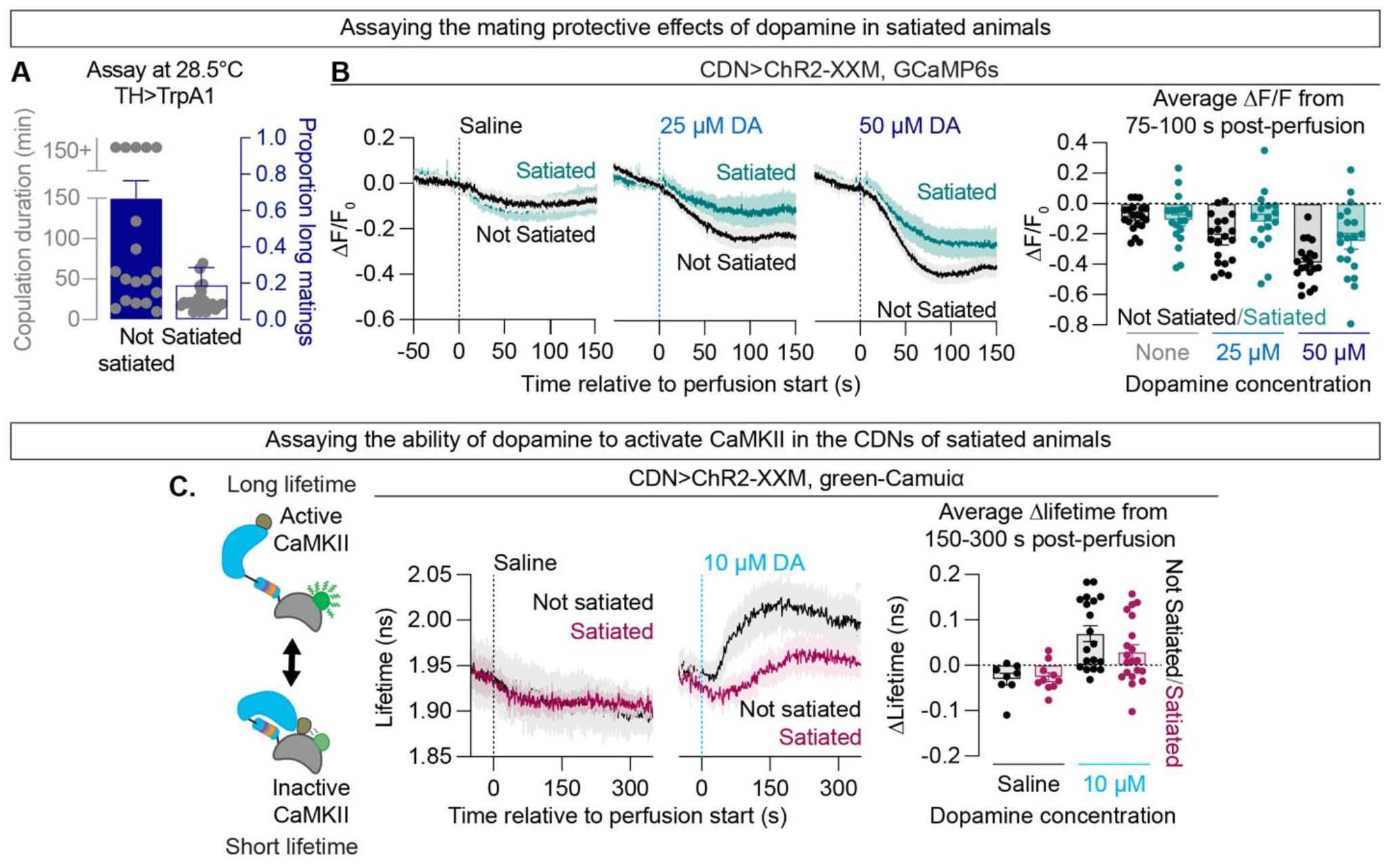
Matings desensitize the CDNs to future dopamine. **A**.Tonic thermogenetic stimulation of dopaminergic neurons extends copulation duration less effectively in satiated males (n=16-21). Grey dots and left y-axis show individual mating durations. Bars and right y-axis show the proportion of matings lasting more than 35 mins ± a 67% credible interval. **B**.Left: Saline controls show no effect of satiety when the CDNs are stimulated by ChR2-XXM and our laser, with calcium dynamics monitored by GCaMP6s (n=19-20). Middle: The CDNs of satiated males are less sensitive to bath application of 25 µM or 50 µM dopamine (n=17-20). Right: summary of the GCaMP data with each dot representing the average change in fluorescence between 75 and 100 s post-perfusion. **C**.Cartoon: The FRET sensor green-Camuiα reports a longer fluorescence lifetime when CaMKII is in its open, active conformation. Left: Satiety has no effect on CaMKII activity in the absence of dopamine (n=8-10). Middle: The CDNs of satiated flies are less sensitive to the CaMKII activity-promoting effects of dopamine application (n=19-20). Trace averages are reported as mean ± SEM. See supplement for individual traces. Right: summary of the green-Camuiα data with each dot representing the average change from baseline in lifetime between 150 and 300 s post-perfusion.

To directly test the idea that prior matings decrease the ability to *receive* the resiliency-promoting dopamine signal, we examined the effectiveness of externally supplied dopamine in our CDN imaging preparations. Consistent with our hypothesis, both calcium and CaMKII readouts showed decreased sensitivity to dopamine in satiated animals (**Figure 3B, C, Supplementary Figure 7H-I**). These data provide a simple explanation for the devaluation of copulation after repeated matings: the ability of the decision-making neurons to detect the motivating dopamine signal is impaired.

### Gαo signaling downstream of the D2R can overcome satiety, but cannot induce it

In Drosophila, the D2R is coupled to Gαo 45,46 (**Figure 4A**) which, when knocked down in the CDNs yielded threat-sensitivity and physiological phenotypes comparable to satiety and loss of D2R function (**Figure 4B, C, Supplementary Figure 8A-C**). We used the light-sensitive mosquito rhodopsin eOPN3^47^ (**Figure 4D**) to optogenetically elevate Gαo signaling in the CDNs, finding an increased resilience to threats (**Supplementary Figure 8D**) similar to the activation of dopaminergic neurons. In contrast to dopaminergic stimulation, which could not rescue the demotivated state induced by prior matings, optogenetic activation of Gαo overcame the effects of prior matings and provided real-time protection from threats (**Figure 4E**). We found that like ChR2-XXM, eOPN3 could be activated by our 2-photon laser set to ∼10mW, allowing us to simultaneously stimulate the CDNs, image calcium dynamics, and activate Gαo. Unlike bath application of dopamine, the ability of Gαo signaling to suppress calcium was not dependent on satiety state (**Figure 4F**). This suggested that matings induce a receptor-level change in the ability of the CDNs to respond to dopaminergic inputs. Consistent with this idea, and in contrast to dopaminergic stimulation, prestimulation of Gαo did not appreciably induce a state of fictive satiety (**Figure 4G**) or otherwise change the motivational dynamics of matings (**Figure 4H**). It therefore appears that the D2 receptor itself must be activated for the pathway to become desensitized and induce the satiety state.

**Figure 4:**
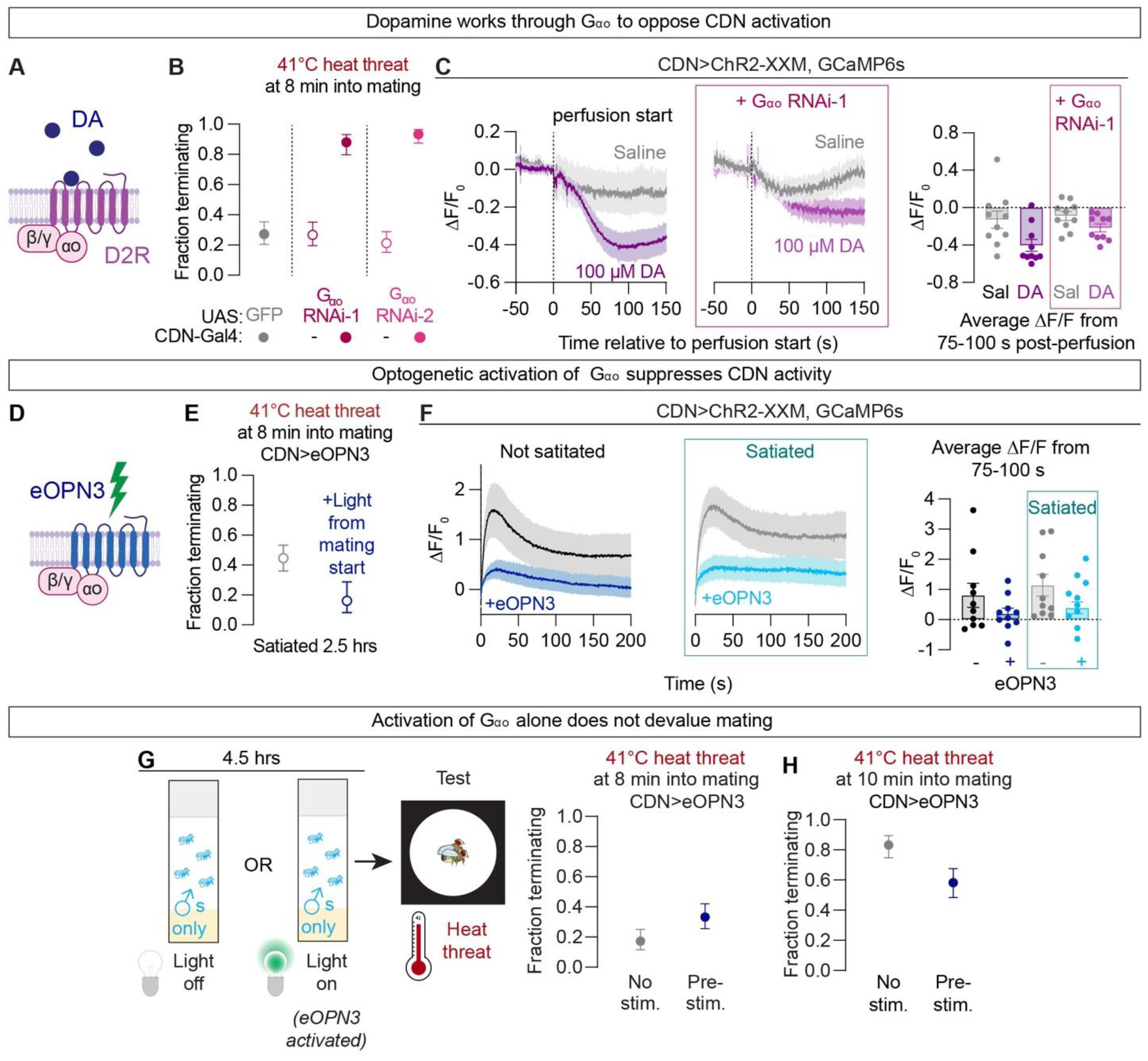
G_αo_ signaling promotes persistence during current matings but cannot induce the devaluation of future matings. A.In Drosophila, the D2R is coupled to G_αo_. B.RNAi-mediated knockdown of G_αo_ in the CDNs using either of two independent RNAi lines increases the fraction of matings that are ended by a heat threat (n=25-28). C.Knockdown of G_αo_ in the CDNs reduced the response to bath applied dopamine (n=10 each). D.The mosquito rhodopsin eOPN3 can be used to optogenetically activate G_αo47_. E.Optogenetic activation of the G_αo_ signaling pathway in the CDNs throughout matings increases threat resiliency in satiated animals (n=13-31). F.Activation of eOPN3 by the 2-photon laser suppresses the calcium response to CDN stimulation (n=10). In calcium imaging experiments with eOPN3 the response was normalized to the first 2 seconds of the recording. Trace averages reported as mean ± SEM. G.4.5 hr pre-activation of G_αo_ signaling in the CDNs produces little, if any, satiety-like increase in the fraction of mating pairs terminating in response to a heat threat during mating (n=29-30). H.Activation of eOPN3 in the CDNs for 4.5 hrs prior to testing does not increase motivation above baseline (n=24 each).

### The classic GPCR inactivation pathway is required for consummatory satiety

Classic GPCR inactivation involves β-arrestin recruitment to activated receptors, preventing future signaling through variety of mechanisms^48^ (**Figure 5A**). Knockdown of the sole Drosophila non-visual β-arrestin (Kurtz)^49,50^ prevented the repetition-induced devaluation of mating (**Figure 5B**) but did not affect the motivational dynamics of unsatiated matings (**Supplementary Figure 9A**). In our imaging preparation, males that had recently mated several times but had β-arrestin knocked down from the CDNs showed no decrease in the ability of dopamine to promote CaMKII activity (**Figure 5C, Supplementary Figure 9B**). This is consistent with the inability of prior matings to alter the response to heat threats in these animals (**Figure 5B**), though deeper investigation points to additional, Kurtz-independent satiety mechanisms (**Supplementary Figure 9C**). These results demonstrate one of the first requirements for this well-studied signaling pathway in controlling natural behavioral dynamics, linking the fields of GPCR inactivation and motivational control. They also argue that without the desensitization pathway acting in dedicated subsets of dopamine-receiving neurons, each bout of a behavior would be treated as though it were the first.

**Figure 5:**
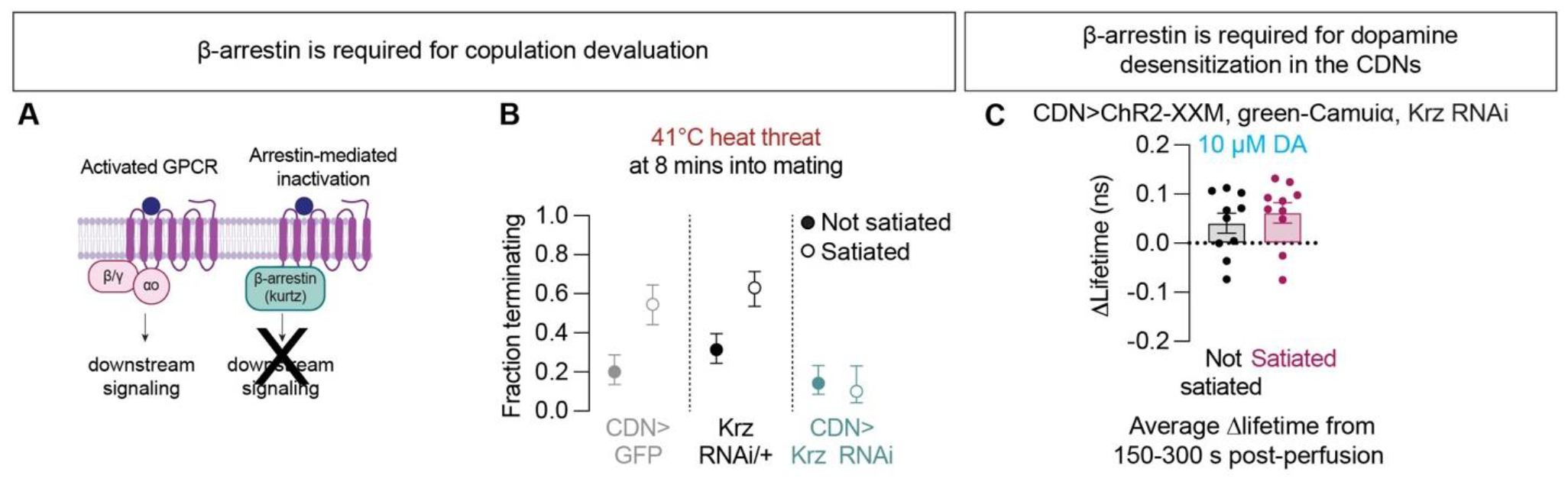
The classical GPCR inactivation pathway is required to implement consummatory satiety. A.β-arrestin (Kurtz) shuts off GPCR signaling by binding to activated GPCRs. B.RNAi knockdown of Kurtz in the CDNs prevents the effects of satiety on termination rate (n=18-21). C.Knocking down Kurtz in the CDNs leaves the CaMKII response to dopamine intact, even after multiple matings (n=10 each).

## Discussion

Studies of motivation have focused largely on neuromodulators and the electrical activity of the neurons that release them, resulting in relatively few molecular-level explanations for motivational state changes^5^. One explanation for this gap has been experimental access to signal-receiving populations: it is often not clear where dopamine and other neuromodulators act to exert their behavior-specific effects. The Drosophila system in general, and reproductive behaviors in particular, allow the level of circuit identification, manipulation, and monitoring required to begin to characterize these mechanisms.

Dopamine is a central signaling molecule for many decisions, including the decision to terminate Drosophila matings when under duress. Earlier analysis of dopamine receptors in this system used copulation duration as a readout^17^. This was misleading because we now know that despite its profound role in decision-making during mating, the duration of unchallenged matings is little affected by loss of dopamine signaling.

We find that the dopaminergic activity that motivated earlier matings desensitizes subsequent matings to future dopamine. If our results are generalizable across species and behaviors, a straightforward prediction is that systemic elevations in dopamine levels in humans would lead to widespread D2R inactivation and consequent devaluation of otherwise rewarding behaviors. This is precisely the case in drug addiction, where broadly acting drugs of abuse cause widespread reduction in D2R availability in the striatum of human addicts^22–25^. This, it seems to us, is an instance of a novel finding in a model organism being effectively pre-validated in humans. Though the desensitization of the D2R is a cornerstone of the dopamine hypothesis of addiction^28,51,52^, we have found no previous suggestion that this mechanism is used to guide natural motivational dynamics at the level of local circuits that govern individual behaviors. Our implication of the D2 receptor desensitization in motivational control, along with molecular, electrical, and behavioral readouts of D2R and CDN function, should enable a deep understanding of the mechanisms that update genetically programmed decision-making circuitry to reflect recent life history.

Studies of addiction in animal models have found reductions in dopamine release^27,51,53^, as well as reception. In our physiology experiments, we supply the dopamine, bypassing changes upstream of the dopamine-receiving cells. The results indicate that the increased sensitivity to threats in satiated males is likely to be due to desensitization at the level of dopamine reception or signal propagation within the CDNs. We suspect that the desensitization occurs at the receptor itself because i) removal of the receptor binding β-arrestin abrogates the effects of satiety and ii) optogenetic stimulation of the downstream signaling molecule Gαo does not induce artificial satiety. None of our experiments rule out a contribution of decreased dopamine release in satiated males, but the inability of dopaminergic stimulation to extend mating in satiated males, together with our ability to rescue motivation by intervening only in the downstream CDNs, argues that the most impactful satiety mechanisms take place in the signal receiving cells.

While dopamine receptor desensitization is likely a broadly conserved satiety mechanism, we know of at least one other behavior for which this is not the case: Drosophila courtship. Male flies use dopamine to motivate courtship through D1-like receptors and display high levels of courtship for days when placed with pheromonally attractive, but behaviorally unreceptive, virgin females^6^ (our observations). This makes it clear that desensitization to dopamine is not unavoidable and implies that desensitization mechanisms can be tuned— or completely inactivated. It may be that, while consummatory behaviors are subject to use-dependent desensitization, appetitive behaviors are not, allowing relentless pursuit of important rewards for hours or longer.

Deeper investigation into this system will likely yield surprises and new insights. The canonical mechanism for GPCR desensitization begins with phosphorylation of active receptors by G-protein Receptor Kinases (GRKs)^54,55^ and the consequent recruitment of β-arrestin^56,57^, often followed by receptor internalization^58,59^. We do not yet know the details of this regulation in our system. No available RNAi lines against GRKs altered the dynamics of copulation when expressed in the CDNs (not shown), but GRK-independent mechanisms of β-arrestin recruitment to GPCRs have been reported^60–63^. Knockdown of β-arrestin blocks desensitization both at the level of the behavioral response to heat threats and the physiological response of the CDNs, but our direct CDN stimulation experiments hint at the existence of β-arrestin-independent copulation satiety mechanisms (**Supplementary Figure 9B**). We do not yet know the ultimate fate of the D2R: whether it is removed from the membrane or inactivated in satiated males through some other mechanism. We also do not understand how the dopamine signal ultimately decreases stimulation-evoked calcium or increases CaMKII levels. Working out these intracellular signaling mechanisms should be possible, either by testing hypotheses gathered from the addiction or GPCR inactivation fields, mining RNAseq data^41^, or from targeted or unbiased screening. The highly conserved mechanisms described in this paper provide a simple solution for a widespread regulatory phenomenon, with clear implications for the motivational dynamics governing a variety of behaviors and decisions. Given the parallels to known mechanisms of addiction, future work on this system may also be of use in understanding the causes and consequences of D2R desensitization underlying a variety pathologies and treatment limitations.

## Acknowledgements

We thank: Stephen Thornquist for UAS-eOPN3; the Rogulja and Crickmore labs for discussions and comments on the manuscript; Ethan Glantz for the design of the behavioral arenas for copulation duration with thermogenetics; the following undergraduates for assistance performing experiments: Kyle Thieringer, Liam Kerrick, Roshinie Persaud, Peter Rifkin, and Annmaria Antony; Maxi Pitsch for assistance analyzing single-cell sequencing data; Ofer Mazor and Pavel Gorelik (Harvard Medical School Neuroinstrumentation Core) for technical advice designing experimental apparatuses; the IDDRC Cellular Imaging Core, funded by NIH P50 HD105351; Thomas Schwarz, Josh Kaplan, Brielle Ferguson, Steven Flavell, Lisa Goodrich, Stephen Zhang and Karina Lezgiyeva for discussions. This work was funded by the NIH (R01NS111441, R01GM134222). LEM was supported by the National Science Foundation Graduate Research Fellowship (DGE 2140743).

## Author Contributions

LEM performed the experiments with early assistance from AKG. LEM and MAC wrote the paper, with input from AKG. All authors designed experiments and analyzed results.

## Methods

### FLY STOCKS

Flies were maintained on conventional cornmeal-agar-molasses medium in narrow vials under 12 hour light/12 hour dark cycles at 25°C and ambient humidity. We did not observe any dependency on time of day on any of the behaviors described here, and all experiments were carried out between ZT0 and ZT12 when the lights would be on.

### Males

Due to the focus of this study, all experimental subjects were adult males. After eclosion, males were anesthetized with CO_2_ for collection then group-housed away from females for at least 3 days before testing. All males used for behavior and imaging experiments were 3-15 days post-eclosion. Flies expressing CsChrimson, GtACR1, ChR2-XXM, or eOPN3 and all corresponding parental controls were fed all-trans retinal for at least 24 hours (Sigma Aldrich #R2500 or Spectrum Chemical R3041) diluted to 50 mM in ethanol on rehydrated potato food (Carolina Bio Supply Formula 4-24 Instant Drosophila Medium, Blue), unless marked as “no retinal”. All retinal vials were kept inside aluminum foil sheaths to prevent degradation of the retinal due to light exposure.

### Females

Virgin females used as partners for termination and copulation duration assays were generated by heat-shocking a UAS-CsChrimson-mVenus stock with a hs-hid transgene integrated on the Y-chromosome (BDSC #55135) in a 37°C water bath for 70 minutes. This stock was selected for mating partners because the females are highly receptive to courtship. Virgin females used in satiety assays were generated by heat-shocking a w^1118^ stock with a hs-hid transgene integrated on the Y-chromosome (BDSC #24638) in a 37°C water bath for 70 minutes. This stock was selected because all experimental males had orange or red eyes, easily distinguishable from this stock’s white eyes when removing the males from satiety assays. Stocks with the hs-hid transgene were maintained in bottles containing conventional cornmeal-agar-molasses media at 19°C. All virgins were group-housed for at least 2-3 days before use.

### Additional notes on fly stocks

No fly used for behavior was anesthetized with CO_2_ within 24 hrs of behavior, all of our flies are aspirated into behavioral chambers from their food vials.

The effects of dopamine stimulation on copulation duration became less consistent with age (data not shown), we suspect due to compensation or decreased transgenic expression. Phenotypes were reliable within the 3-15 day age range used in this study.

All RNAi/+ controls were produced by crossing the RNAi line to Canton-S virgins. All TrpA1/+ controls were produced by crossing UAS-TrpA1 virgins to Canton-S males.

The D2R^Δ1^ CDN>UAS.D2R rescue experiment flies used the following mating scheme:

1)To put D2R^Δ1^ with CDN-Gal4, BDSC stock #78795 was double balanced in the w^1118^ background then crossed to add NP2719-Gal4, Repo-Gal80 to the second chromosome (aka CDN-Gal4). The resulting stock was D2R^Δ1^; CDN-Gal4/CyO; TM2 or 6B/+. The floating balancer on the third chromosome was selected against when collecting males for behavior.

2)To isolate UAS-D2R.S on the third chromosome, BDSC stock #78798 was balanced in the w^1118^ background (removing D2R^Δ2^ from the X chromosome), then crossed to a w^1118^ stock with balancers only on the third chromosome. The resulting genotype was +; If or CyO/+; UAS-D2R.S/6B. The floating balancer on the second chromosome was selected against when collecting males for behavior.

3)For the CDN>UAS-D2R control group males of D2R^Δ1^; CDN-Gal4/CyO; TM2/+ were crossed to virgins of +; If or CyO/+; UAS-D2R.S/6B. For the experimental group, virgins of D2R^Δ1^; CDN-Gal4/CyO; TM2/+ were crossed to males of +; If or CyO/+; UAS-D2R.S/6B.

These stocks are available upon request.

We found eOPN3 to have dark activity that worsens with time on retinal, and so tested these animals within 24-28 hrs of retinal feeding for behavioral experiments.

For all imaging experiments with CDN>ChR2-XXM, GCaMP6s or green-camuiα and an RNAi, a stable stock with Dicer2, CDN-Gal4, Repo-Gal80, and the RNAi was created first then crossed to ChR2-XXM and the requisite imaging tools.

### SATIETY ASSAYS

In the satiety assay, individual male flies were aspirated into a vial containing conventional cornmeal-agar-molasses medium and ∼15 virgin w^1118^ females. Vials containing 15 virgin females were prepared at least 2-3 days before use for satiety assays. In general, we used females that appeared healthy – climbing up the walls and moving around. Satiety vials are usually used within 3-5 days of preparation, and never used more than 20 days after preparation, resulting in an age range of 3-10 days old for the females used in the vast majority of experiments.

Satiety assays are run in the lab at ambient temperature (∼23°C), humidity, and lighting unless otherwise noted. We did not monitor the number of matings that occur in each satiety assay. In our experience^6,19,34^ it is extremely rare for an animal not to mate multiple times in a satiety assay. For all experiments, control genotypes were satiated simultaneously with experimental genotypes.

Males are always aspirated into and out of satiety assays. If a male is still mating at the end of the assay, the mating pair is broken up by repeated aspiration so he can be removed. All males were used for behavioral testing within one hour of removal from the satiety assay.

In general, 2.5 hr satiety assays were used for subsequent behavioral testing because we needed the male to mate again, and courtship is more strongly reduced after 4.5 hours in a satiety assay. The one exception is **Figure 1C**, where a 4.5 hr satiety assay was used to be able to compare the results to the 4.5 hrs thermogenetic pre-stimulation experiments. For imaging experiments where the males are not required to mate again afterwards, we were able to more fully satiate males using a 4.5 hr satiety assay.

*Satiety recovery experiments*

For any experiments with a recovery period following a satiety assay (e.g., **Figure 1B**), males were satiated as described above then aspirated into a new vial at the conclusion of the satiety assay. Males were housed in groups of 10-12 in standard conditions before testing.

## BEHAVIORAL EXPERIMENTS

### Evaluation of mating

#### Automated

For copulation duration experiments not involving thermogenetics or optogenetics, males and females were loaded into 32-well arenas (https://github.com/CrickmoreRoguljaLabs/FlyBehaviorChambers, see also Boutros, Miner, et. al. 2017) recorded from a height of ∼9” using a Canon camera (VIXIA HF R600) with videos saved on an SD card. The recordings were then processed using a custom MATLAB code (also at https://github.com/CrickmoreRoguljaLabs/FlyKnight). For additional details see Thornquist et al., 2020. Results from the MATLAB code were spot checked by an experimenter for accuracy, especially in the case of extremely short (to check that a true mating occurred) or extremely long (to confirm the pair is not stuck) matings. If a pair began mating before the start of the video, as happens occasionally when moving the plate from the loading area to the cameras, that data point was excluded.

#### Manual

Termination assays were evaluated in real time by the experimenter, with video recordings as backup. Manually scored copulation duration assays (i.e., the TH>TrpA1 experiment in **Figure 3A**) were recorded and scored *post hoc* by an experimenter to an accuracy of ∼10s using VLC media player.

A pair was scored as mating when they adopted a stereotyped mating posture for at least 30 seconds. This posture consists of the male mounting the female and propping himself up on her abdomen using his forelegs, while curling his abdomen and keeping the genitalia in contact (**Supplementary Figure 10A**). The posture is starkly different from anything exhibited during other naturalistic behaviors and easily recognizable by a trained experimenter. On rare occasions the male will exhibit a “stuck” posture while attempting to terminate the mating. When stuck, the male dismounts the female, orients himself away from her and attempts to free himself but cannot decouple their genitalia (**Supplementary Figure 10B**). In the rare cases when we see a male become stuck in response to a threat, we exclude that mating. The stuck phenotype is slightly more frequent in extremely long matings (>1 hr), possibly due to the hardening of released seminal fluids that adhere the flies together. In this case, the end of mating is scored as the onset of the stuck posture.

TH>TrpA1 stimulation experiments often caused a rearing-like posture that does not meet our criteria for being stuck. In the these experiments, the male often rocked back from the female, but did not orient away from her or otherwise attempt to disengage (**Supplementary Figure 10C**). The male will often move between this rearing position and the stereotypical mating posture, whereas a stuck male will never return to the stereotypical mating posture. Males in the rearing posture are therefore scored as mating.

#### Termination of mating

For termination assays, matings must terminate within 1 minute after the end of the optogenetic stimulation or heat threat, to be scored as “terminated”. In experiments where both optogenetic silencing and heat threats are use, matings must terminate within 1 minute of the end of the heat threat (e.g. **Figure 2A**). It is very rare to see truncations that occur more than 1 minute after a stimulus has ended.

For copulation duration assays, the end of mating was scored when the mating pairs separate or become stuck (see above). For genotypes with extremely long copulations (e.g., TH>TrpA1), copulation durations were capped at 150 minutes, after which the data points were pooled (as in **Figure 3A**). This cutoff was decided after pilot experiments showed that flies mating that length of time were highly likely to continue mating for many more hours, often until death. When a mating pair dies in the act of mating, they often remain in the stereotyped mating posture. This made it difficult to reliably determine the point of death from the videos to use as the end of mating.

### Additional notes about behavior

During this study we found a previously unreported deficit in the ability of D2R whole animal mutants to mate^6,17^, with less than 5% of males mating (data not shown). This is unlikely to be due to expression of the D2R in the CDNs, as CDN>D2R-RNAi flies did not show any deficits in mating ability.

Only flies that were mating when the stimulus began were included in the termination experiments. This accounts for almost all experimental flies, but on the rare occasion that a mating terminated naturally before stimulus delivery, that pair was excluded.

#### Calculating the proportion of long matings

Matings were classified as long if they were over 35 mins. This length is almost never observed in control genotypes in our experience.

### Behavioral arenas for copulation duration with thermogenetics

One male and one female were placed in a cylindrical acrylic well (10 mm diameter by 3 mm height) within a 32-well plate. The floor of the well consisted of diffuser film (Inventables, 23114-01) separating the chamber from a water reservoir that sits beneath all wells. The temperature of the chambers is controlled by constantly pumping water from a water bath through this reservoir. The temperature of the water bath was set so that a thermocouple touching the floor of the well read 28.5°C for stimulation experiments or 25°C for controls. Videos were recorded with a Canon camera (VIXIA HF R600) and saved on an SD card for an experimenter to score afterwards. Schematics are available at https://github.com/CrickmoreRoguljaLabs/ThermogeneticsBox.

### Behavioral arenas for termination assays

#### General setup for termination assays

For more information on the behavioral areas we used, see Thornquist et al., 2020 and https://github.com/CrickmoreRoguljaLabs/Waterworks. Briefly, one male and one female were placed in a cylindrical acrylic well (86 mm diameter by 3 mm height) with a diffuser film (Inventables, 23114-01) floor. Each well sits above a¼ inch thick water reservoir, whose source can be controlled by the experimenter to deliver heat threats. Threat strength was measured by inserting a thermocouple into the behavioral well touching the floor.

#### Termination assays with optogenetics or a blue light stressor

For CsChrimson experiments: One male and one virgin female fly were placed in each 86 mm diameter 1/8” thick acrylic well sitting 4” above 655-675 nm LEDs (Luxeon Rebel, Deep Red, L1C1-DRD1, formerly LXM3-PD01-0350) driven using a 1000 mA constant current driver (LuxDrive BuckPuck, 03021-D-E-1000) and passed through frosted collimating optics (Carclo #10124). This spot of light was scattered using a thin diffuser film (Inventables, 23114-01) under the wells to ensure a uniform light intensity of ∼0.1 mW/mm^2^. This film also forms the floor of the behavioral arena. The LEDs were controlled using an Arduino Mega2560 (Adafruit) running a custom script, which itself was controlled by a Raspberry Pi (either 2 or 3, running Raspbian, a Debian variant). Flies were observed by recording from above using the Raspberry Pi with a Raspberry Pi NoIR camera (Adafruit) and infrared illumination from below using IR LED arrays (Crazy Cart 48-LED CCTV IR Infrared Night Vision Illuminator reflected off the bottom of the box) while streaming the video to a computer for observation.

For GtACR1, ChRmine, and eOPN3 experiments: The set-up was as above except using a 520-540 nm LED (Luxeon Rebel, Green, L1C1-GRN1, formerly LXML-PM01-0100) driven using a 700 mA current driver (LuxDrive BuckPuck, 03021-D-E-700), and a pulse-width modulated signal to set the time-average intensity to ∼10 µW/mm^2^ unless otherwise noted.

For blue light stressor experiments: The set-up was as above except using a 465-485 nm LED (Luxeon Rebel, Blue, L1C1-BLU1, formerly LXML-PB01-0040) driven using 1000 mA constant current drivers (LuxDrive BuckPuck, 03021-D-E-1000) providing an intensity of ∼167 µW/mm^2^.

For all experiments with optogenetic stimulation beginning at mating start, the LED was turned on at the start of mating and off at the end of the experiment. For all experiments with optogenetic stimulation surrounding a heat threat, the LED was turned on 30s before the beginning of the heat threat and turned off 30s after cessation of the threat.

#### Thermogenetic stimulation during termination assays

For experiments where there is thermogenetic stimulation and heat threats (i.e., **Supplementary Figure 3E**), two water baths were connected to our behavior box (see Thornquist., 2020 for a diagram). This allowed for warm water to flow throughout the assay except during the heat threat.

### Stimulation and silencing outside of a termination assay

#### Thermogenetic pre-stimulation (including ejaculatory bulb depletion)

For thermogenetic pre-stimulation, males were loaded into vials and then placed in either a 23°C, 28.5°C, or 30°C incubator with the lights on and ∼50% humidity for 4.5 hrs. In general, stimulation was performed at 30°C unless the flies showed, or had previously been reported to show, seizure-like behaviors (TH-GAL4^6^, Crz-GAL4).

#### Optogenetic pre-stimulation

For eOPN3 experiments: a lightbox was created by affixing 10 6-LED arrays (FXC 6 LED Clearance Truck Bus Trailer Side Marker Indicators 12V, Green) in parallel ∼1/2” apart to a 12.5” x 7” piece of 1/8” black acrylic forming the base of a box with 1” high sides made of 1/8” black acrylic. The top of the lightbox was made from a 1/16” piece of clear acrylic topped with diffuser paper (Diffusion Gels Filter Sheet Kit, #No.3 Light Frost, 54% transparency). Vials are laid sideways on the box in line with each LED array, resulting in ∼8.12 mW/mm^2^ light intensity. Schematics are available at https://github.com/CrickmoreRoguljaLabs/GreenLightpads.

#### Silencing during a satiety assay

The apparatus described above was also used for all GtACR1 experiments that involved silencing during a satiety assay. For TH>GtACR1 experiments, an experimenter continually observed satiety assays. When a male began a mating, his vial was moved to the lightbox and laid on its side until the mating ended. For Crz>GtACR1 (for ejaculatory bulb assessment) experiments, the vials were left on the box for the entire, unsupervised satiety assay.

#### TH silencing experiments when untethered to mating

The set up was the same as described above for TH>GtACR1 experiments in the satiety assay. Here, after a 2.5 hr satiety assay, males were aspirated from their satiety vial into a conventional cornmeal-agar-molasses food vial with the other males from that session (∼6-12 males). Vials were then placed on top of the lightbox for five 20-min light treatments with 20-min light-off breaks in between.

### Scoring conventional satiety in the satiety assay

Males were loaded into satiety vials and lined up on a bench top in the lab away from activity so the experimenter could observe behavior without disturbing the vials. The presence or absence of mating behaviors (courtship and copulation) was scored manually every 30 min for 4.5 hours, as in groups of 6 vials. A within-group percentage was generated for each time point.

## IMAGING EXPERIMENTS

### Saline preparation

External saline (Wilson and Laurent, 2005^64^) was prepared for all imaging experiments. This saline is composed of 103 mM NaCl, 3 mM KCl, 5 mM TES, 8 mM trehalose, 10 mM glucose, 26 mM NaHCO_3_, 1 mM NaH_2_PO_4_, 3 mM MgCl_2_, and 1.5 mM CaCl_2_ with a pH ∼7.3 and 270-275 mOsm.

### Dopamine preparation

10 mM dopamine-HCl (TOCRIS #62-31-7) in water was prepared in small batches to a then stored at -20°C in 1000 µL alliquots in black Eppendorf tubes (Fisher Scientific #15386548) until ready for use. On the day of the experiment, an aliquot was defrosted, and the dopamine further diluted in 20 mL of saline to the desired concentration for each experiment. This dilution was performed immediately prior to each recording in a new 50 mL Falcon tube (Corning #352098).

### Ejaculatory Bulb Volume Scoring

Males were immediately dissected following treatment in external saline. Ejaculatory bulbs were mounted on to the base of a 35 x 10 mm petri dish (VWR #10799-192). The petri dish was placed on a piece of black acrylic (to create a contrasting background) and photographed using a dissection microscope (Nikon SMZ745T) mounted with an Excelis HDMI camera (ACCU-SCOPE #AU-600-HD) captured with CaptaVision+ software. Dissection, mounting, and image acquisition were performed by LEM. LEM then blinded the images and gave them to MAC, who scored them on a scale from 1 (empty) to 4 (full) as previously described^6,17^.

### Immunostaining and fixed microscopy

The nervous system of the fly was removed in external saline and immediately fixed in 4% paraformaldehyde in 0.3 % PBST at room temperature for 20 min. After washing with 0.3% PBST three times (20 min each), the nervous system was incubated for 2 hrs with 2% normal donkey serum (Jackson ImmunoResearch #AB_2337258) in 0.1% PBST at room temperature. The nervous system was then incubated with primary antibodies (diluted in the same blocking solution) at 4°C for 3 days. Following a 0.3% PBST wash (three times for 20 mins), the nervous system was incubated with secondary antibodies (diluted in bocking solution) at 4°C for 3 days. After another 0.3% PBST wash (three times for 20 mins), the brains were mounted with ProLong Diamond Antifade Mountant (ThermoFisher Invitrogen #P36961) on glass slides (Azer Scientific #EMS200WS) with micro cover glass No.1 (VWR #48393-026) using standard procedures. Confocal sections were acquired using a Leica Stellaris 5 laser scanning confocal microscope at 1024 x 1024 pixels. For images of the entire ventral nervous system (VNS; **Supplementary Figure 3B**) images were collected in slices that span the entire VNS using a 20x immersion objective (HC PL APO CS2 20X/0.75) at a 3 µM interval (approx. 45 slices). For images of the abdominal ganglion (**Figure 2B**) images were collected in slices that span the entire abdominal ganglion using a 63x glycerol objective (HC PL APO CS2 63X/1.30) at a 1 µM interval (47 slices total). All shown images are maximum projections of image stacks obtained in FIJI (Schindelin et al. 2012).

### 2-photon Imaging: intensity and FLIM experiments

With the exception of **Supplementary Figures 6B and 7B** (see *Baseline calcium imaging* below), all GCaMP intensity and FLIM experiments were based on the experiments in (Gautham et al., in revision), as summarized below.

All satiated males used in imaging experiments had been in the satiety assay for 4.5 hours and were used within a couple of hours afterward.

#### Microscope

We used a modified Thorlabs Bergamo II with a Coherent Chameleon Vision II Ti:Sapphire laser emitting a 920 nm beam to excite samples. Light was collected through a 16x water objective (Nikon, 0.8 NA, #N16XLWD-PF) immersed in the petri dish with chilled saline and emission was detected using cooled Hamamatsu H7422P-40 GaAsP photomultiplier tubes. The PMT signal was amplified using fast PMT amplifiers (Beker-Hickl #HFAC-26) then passed to a PicoQuant TimeHarp 260 photon counting board synchronized to the laser emission by a photodiode (Thorlabs #DET110A2) inverted using a fast inverter (Becker-Hickl #A-PPI-D). Custom software (FLIMage, Florida Lifetime Imaging, Version 2.0.21, available at https://github.com/ryoheiyasuda/FLIMage_public) was used to acquire the TimeHarp signal and control the microscope.

#### Perfusion system

Dopamine or saline was perfused into the petri dish containing the sample during imaging using the setup described in Gautham et al., in revision. Briefly, a 50 mL Falcon tube (Corning #352098) containing 20 mL of the solution to be perfused was elevated ∼21” above the sample. The solution was connected to the sample by a 1’ of 1/8” tubing (McMaster-Carr #5648K74), followed by a three-way stopcock (Smiths Medical #MX5311L), followed by a 1.5’ of 1/8” tubing which was then connected to 1’ of 1/32” tubing (McMaster-Carr #5233K91) via a 1000 µL pipette tip (VWR #76322-154). Connections were sealed with commercially available hot glue where needed. To the other end of the 1/32” tubing we attached a 10 µL pipette tip (VWR #53509-135). This tip was held in place by a stand and could be interested into the petri dish containing each sample by the experimenter. To control the timing of perfusion we attached a 20 mL leur lock syringe (LiteTouch) to the stopcock.

To maintain a constant level of liquid in the petri dish while imaging, old solution was removed by a similar tubing system, with a 10 µL pipette tip connected to 1/32” tubing, connected to 1/8” tubing, terminating into a 250 mL bolt neck flask with upper tubulation via a rubber stopper with a hole inserted into the mouth of the flask. Tubing was used to connect the flask to a vacuum system.

#### Dissection

All imaging experiments use an *ex vivo* preparation of the ventral nervous system. For GCaMP6s experiments, flies were anesthetized with CO_2_, dipped in 100% ethanol, then dissected in chilled saline. For green-Camuiα experiments, flies were instead anesthetized with ice then followed the same procedure, as we found CO_2_ anesthesia sometimes affected baseline green-Camuiα activity and made it unresponsive to stimulation. The ventral nervous system was mounted dorsal-side-down in a 35 x 10 mm petri dish (VWR #10799-192) with 4 mL of chilled saline.

#### Region of interest selection

The CDNs have bilateral axonal projections within the abdominal ganglion that form distinct half-circle projection patterns (see Thornquist & Crickmore, 2020, Gautham et al., in revision). We use a region of interest encompassing the axons on either side of the abdominal ganglion, depending on the strength of fluorescence and clarity of the morphology.

The dopamine neurons labeled by the TH-Gal4 driver have expression throughout the abdominal ganglion. We chose an ROI that showed the strongest fluorescence and most consistent morphology between samples, which was a circular region in the center at the base of the abdominal ganglion containing dendrites. This region can also be identified by TH>Denmark staining.

For GCaMP experiments the region of interest and background selection were chosen selected within FLIMage software at the start of each experiment. For FLIM experiments the region of interest was chosen within the GUI for our custom software, see below.

#### Acquisition

Images were acquired at 64 x 64 pixels at a frame rate of ∼7.8 Hz (1 frame every 0.128 seconds). For GCaMP experiments without perfusion (**Figure 5F**) we used a 200s recording (1570 frames). For all GCaMP experiments with perfusion, we used 300s recordings (2350 frames) and began perfusion at 150s into the recording allowing sufficient time for the activity to stabilize (**Supplementary Figure 5B**). We found that CaMKII activity takes longer to change in response to dopamine than calcium dynamics, therefore for all green-Camuiα perfusion experiments we used a 450s recording (3910 frames), beginning the perfusion at 150s into the recording to match our GCaMP experiments. We did not perform any averaging at the acquisition stage.

#### Perfusion during imaging

Prior to the start of imaging for any experiment where perfusion was performed, 20 mL of a fresh dilution of dopamine or an aliquot of saline was prepared in a 50 mL Falcon tube. The Falcon tube was affixed with a clamp to the top the perfusion system and connected to the perfusion system tubing. A clean syringe was attached to the stopcock and 5 mL of solution loaded. During the experiment, starting at 150s, the solution was carefully pushed from the syringe through the tubing leading to the sample and the stopcock subsequently turned to allow free flow from the 50 mL tube to the sample at a rate of ∼4 mL/min. If the perfusion caused the sample to be displaced or detached from the petri dish, the recording was stopped, the data excluded, and the sample thrown out. Samples were never reused once a recording began.

#### Determination of dopamine concentration

To determine the effective concentration of dopamine we referenced Gautham et al., in revision and performed pilot experiments. We found 100 µM DA to be a high-end effective dosage for GCaMP experiments. We found that 10 µM DA was effective for Camuiα experiments and that 25 µM and 50 µM concentrations did not have any appreciably stronger effects in naïve or satiated animals.

#### Optogenetic stimulation during imaging

We recently discovered (Gautham et al., in revision) that our 2-photon laser set to 920 nm and operating at sufficient power could be used to simultaneously activate certain optogenetic tools, while also stimulating GCaMPs or green-Camuiα. We take this approach throughout the paper to activate ChR2-XXM and/or eOPN3. To ensure a consistent activation of these tools over time, before each imaging session we measured the power of the laser at the sample using a Coherent FieldMate Laser Power Meter 1098948 and set the % laser power in the FLIMage software so that our Laser Power Meter read between 10 and 11 mW (we aim for 10.5 but the power meter is a bit noisy). The actual value of the % laser power varied across days but was approximately 30 ± 2% across experiments.

For **Supplementary Figure 7C**, ChR2-XXM was used instead to acutely stimulate the dopaminergic neurons. In this case, the experimenter took a couple baseline frames (data not shown) adjusting the power until finding one that causes approximately

200 photons to be detected in the ROI. This typically corresponded to a laser power around 3-4 mW, or 12% power in the FLIMage software. Optogenetic stimulation was performed by excitation with a blue (Thorlabs #M470L4) mounted LED through a liquid light guide (Thorlabs #LLG5-8H) resulting in an incident intensity at the sample of approximately 0.2 mW/mm^2^. Our 2-photon laser and PMTs are shuttered during stimulation using an Arduino, which closed the shutter 100 milliseconds before and opened the shutter 100 milliseconds after the stimulation to protect the PMTs.

#### GCaMP experiment analysis

GCaMP experiments were analyzed using FLIMage, summing all detected photons within a pixel/frame together regardless of arrival time. To then calculate normalized fluorescence intensity values, .csv files containing the background subtracted fluorescence intensity (reported as photon count per frame) were exported from the FLIMage software and a custom MATLAB script (available at https://github.com/CrickmoreRoguljaLabs/GCaMPfromFLIMage ) was used to calculate ΔF/F values. The median of the baseline was used to calculate F_°_. For perfusion experiments, the baseline used was the 10 seconds immediately prior to the start of perfusion (140-150 seconds into the recording). For eOPN3 experiments, the baseline used was the first 2 seconds of the recording. For acute optogenetic stimulation experiments, the baseline was the 30 seconds immediately prior to the start of the blue light pulse. On occasion, we observed noise in our imaging system that caused an artifact producing photon values orders of magnitude above baseline, visible as a straight line across the screen. We excluded the few frames with this artifact manually.

#### FLIM experiment analysis

For FLIM experiments we use FLIMage, to record the arrival time of each photon emitted relative to the time of each excitation pulse delivered by our laser (operating at 80.310 MHz) for each pixel within the ROI. We did not average across frames in the acquisition phase. To then calculate the fluorescent lifetime values, .FLIM files were exported from the FLIMage software and analyzed using custom software described in detail in Gautham et al., in revision and available at https://github.com/CrickmoreRoguljaLabs/flim-analysis. In this software we averaged every 10 consecutive frames together. On occasion, we observed noise in our imaging system that caused an artifact producing photon values orders of magnitude above baseline, visible as a straight line across the screen. We excluded the few frames with this artifact manually.

### Baseline calcium imaging

For **Supplementary Figures 6B and 7B**, we used a Zeiss LSM 710 Multiphoton Confocal with a Mai-Tai laser set to a wavelength of 960 nm in order to excite both GCaMP6s and tdTomato and analyzed the data as previously described in Zhang 2016. Briefly, we used a W Plan-Apochromat 20x/1.0 DIC VIS (IR) M27 75 mm objective to take ∼ 40 sections spanning the abdominal ganglion at 3 µM intervals and then acquired maximum projections of each stack in FIJI. Image files were blinded and randomized using a custom MATLAB code (available at https://github.com/CrickmoreRoguljaLabs/GCaMP-tdt_Analysis), then an experimenter used the tdTomato channel to selected an ROI encompassing the entire abdominal ganglion and background ROI. The MATLAB code was then used to compute first the average pixel fluorescence in each ROI in each channel, and then the normalized fluorescence as the (GCaMP_ROI_ - GCaMP_background_)/(tdTomato_ROI_ – tdTomato_background_).

## ADDITITIONAL ANALYSIS

### Analysis of available single-cell sequencing data

We used a recently published single-cell sequencing atlas^41^ to confirm the expression of the D2R in putative CDNs. We defined putative CDNs by their expression of CG17646, dsx, and Gad1^17^. 100% of cells defined in this way show detectible expression of the D2R (data not shown).

## QUANTIFICATION AND STATISTICAL ANALYSIS

### Credible intervals for proportions

To set the error bars for proportion of matings terminating in response to a challenge, as in Thornquist et al., 2020, we use a 68% credible interval surrounding the mean calculated from the non-informative Jeffery’s prior when reporting error to approximate the SEM metric, which is the 68% credible interval on the mean under a uniform prior. These calculations were performed using custom code (https://github.com/CrickmoreRoguljaLabs/UpdatedJeffi).

### Hypothesis testing on proportions

As in Thornquist et al., 2020, we use the non-parametric Fisher’s exact test to test the hypothesis that two sample proportions were drawn from the same Bernoulli process. We correct *post hoc* for multiple comparisons using the Bonferroni correction: 0.5/n where n is the number of unique comparisons made.

### Standard error of the mean for continuous (non-binary) data

We use the standard SEM metric to estimate variability in sample means when the data are continuous (e.g., copulation duration and imaging).

### Hypothesis testing on continuous (non-binary) data

We use the non-parametric Mann Whitney-U test on rank sums to test for differences in distributions of copulation durations. We correct *post hoc* for multiple comparisons using the Bonferroni correction: 0.5/n where n is the number of unique comparisons made.

For imaging experiments, to be able to compare between groups we selected a time window that represented when dopamine perfusion had just reached maximum effectiveness and averaged the values for all the frames in that window for each animal to create the summary figures. For GCaMP experiments this was 75 to 100 s after the start of perfusion. For green-Camuiα experiments this was 150 to 300 s after the start of perfusion. We then performed the Mann Whitney U-test on these distributions with the Bonferroni correction described above.

