## Supplementary Figures and Tables for "Local desensitization to dopamine devalues recurring behavior"

#### Supplementary Figure 1

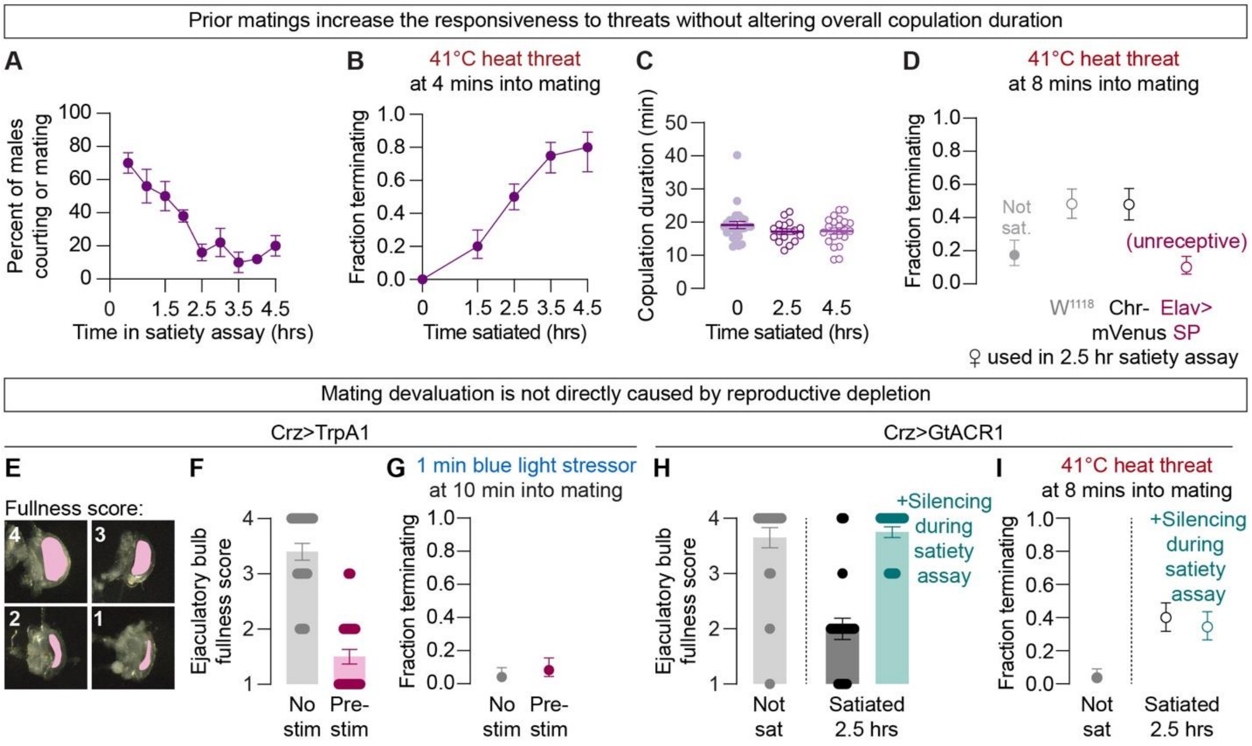

##### Supplementary Figure 1: Consummatory satiety is a function of time spent mating

- **A.** The percentage of males engaged in mating behaviors – either courtship or copulation – declines as they spend more time mating *ad libitum* with females in the satiety assay (n=36, 6 groups of 6).
- **B.** The fraction of males terminating matings in response to the same heat threat increases with time spent in the satiety assay (n=10-38).
- **C.** Copulation duration does not change with satiety (n=16-25).
- **D.** Effectiveness of the satiety assay is independent of the type of female used, so long as the females are receptive. Spending time with unreceptive females does not induce consummatory satiety (n=25-29).
- **E.** Representative images of ejaculatory bulb scores. The lumen that stores sperm and seminal fluid is pseudocolored pink.
- **F.** Thermogenetic stimulation of Corazonin neurons for 4.5 hours causes ejaculation and reduces the ejaculatory bulb volume (n=24-25).
- **G.** Unlike stimulation of the dopamine neurons in **Figure 1C**, 4.5 hours of thermogenetic pre-stimulation of the Corazonin neurons does not induce an increase in the fraction of mating pairs terminating in response to a heat threat (n=24-25).
- **H.** Silencing the male Corazonin neurons throughout a satiety assay prevents sperm transfer from occurring during mating, shown by a still full ejaculatory bulb after the assay (n=20 each). Note that matings with the male Crz neurons silenced are extended to ~90 mins each.
- **I.** Consummatory satiety is still observed when ejaculation was prevented during prior matings. Note that the silencing was turned off during the final, challenged mating (n=27-30).

#### Supplementary Figure 2

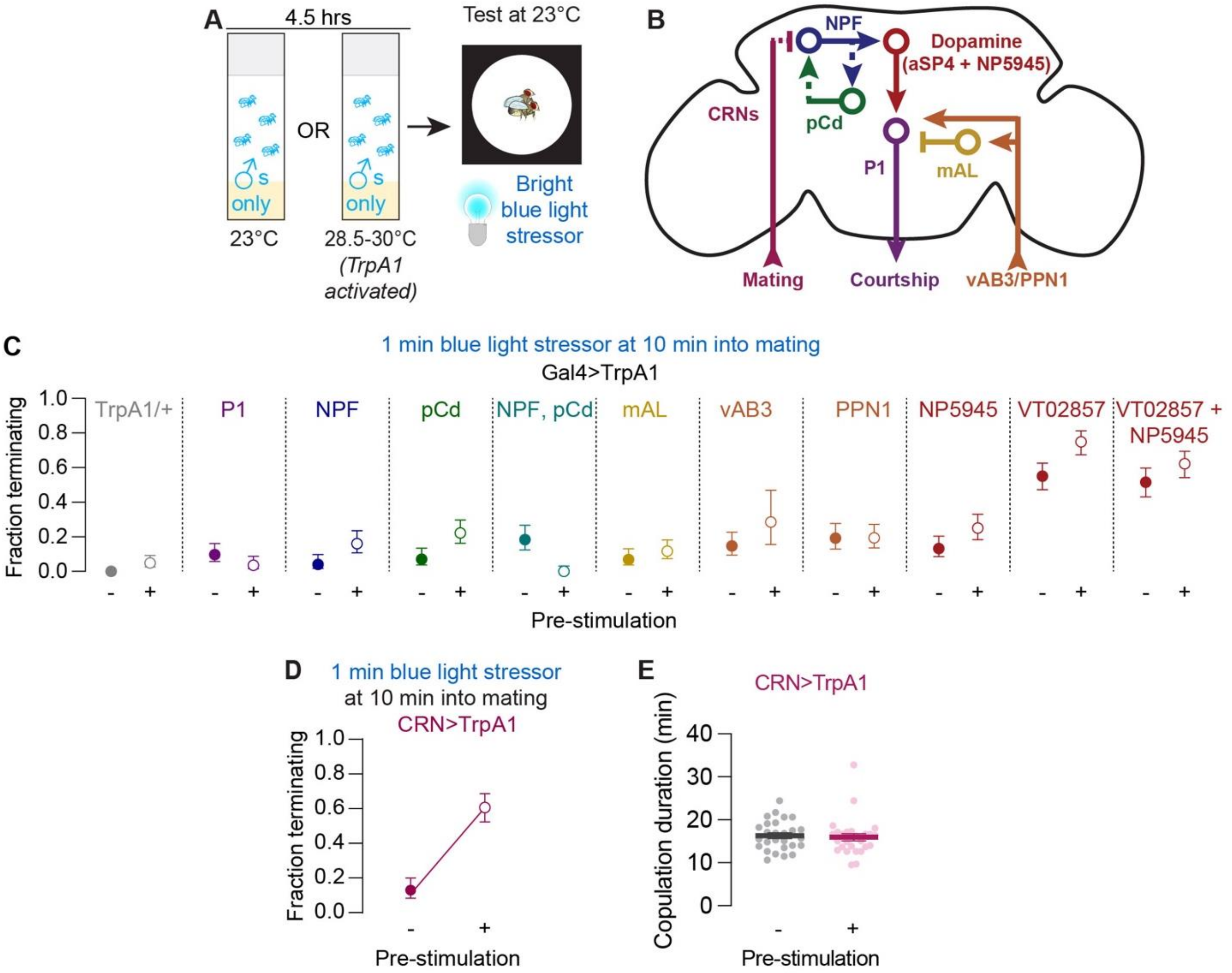

**Figure S2: Testing courtship circuit elements for potential roles in consummatory satiety**

**A.** Males are group-housed in standard food vials and exposed to either warmth to activate TrpA1 or left at room temperature for 4.5 hrs, then their motivation is assessed by testing the probability that they terminate the mating in response to a bright blue light stressor in a subsequent mating. See Methods for explanation of stimulation temperatures.

**B.** Model: known circuit elements controlling appetitive satiety and courtship behavior.

**C.** Pre-stimulation of known courtship circuitry neurons does not induce increased mating termination in response to a pulse of blue light (n=7-41). The high baseline in VT02857 precluded us from testing this Gal4 line, which labels the sexually dimorphic brain dopamine neuron called asp4[64]. We did not follow up because the results in **Figure 1D** indicate that no brain dopamine neurons are important for consummatory satiety.

**D.** Pre-stimulation of the Copulation Reporting Neurons (CRNs; R42G02-Gal4), a putative sensory neuron population that projects to the genitalia and has been reported to detect mating[19], induces increased termination in response to a pulse of blue light (n=31-33).

**E.** Pre-stimulation of the CRNs does not alter copulation duration (n=27 each).

#### Supplementary Figure 3

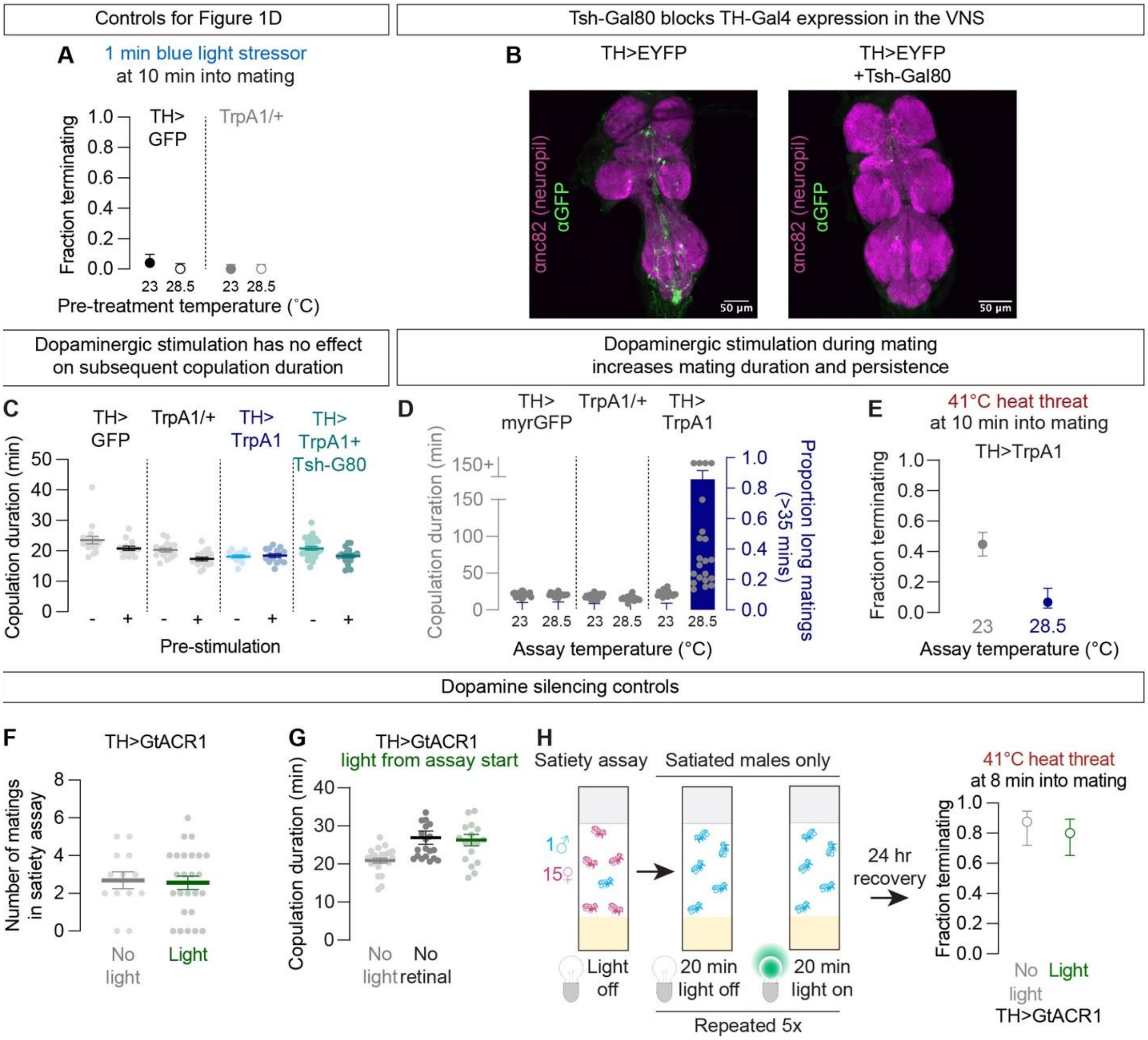

**Figure S3: Dopaminergic release during mating is transiently protective while inducing long-term deprioritization of future copulations**

- **A.** Parental controls for Figure 1D, exposure to the heat treatment used to thermogenetically activate TrpA1 does not itself induce a devaluation of mating (n=25-33).
- **B.** Immunostaining of the ventral nervous system of male Drosophila expressing EYFP without (left) and with (right) the addition of Tsh-Gal80 transgene. Neuropil is in magenta.
- **C.** Pre-stimulation of dopamine neurons does not change copulation duration (n=14-27).
- **D.** Sustained stimulation of dopamine neurons during mating extends copulation duration (n=16-22).
- **E.** Stimulation of dopamine neurons reduces the probability of the mating terminating in response to a heat threat (n=15-38).
- **F.** Electrical silencing of dopamine neurons during mating in a satiety assay does not alter the number of times males mate in the assay (n=13-25).
- **G.** Sustained silencing of dopamine neurons does not alter copulation duration (n=18-23).
- **H.** Silencing dopamine neurons after a satiety assay is not sufficient to counteract the effects of satiety on motivation (n=8-10).

#### Supplementary Figure 4

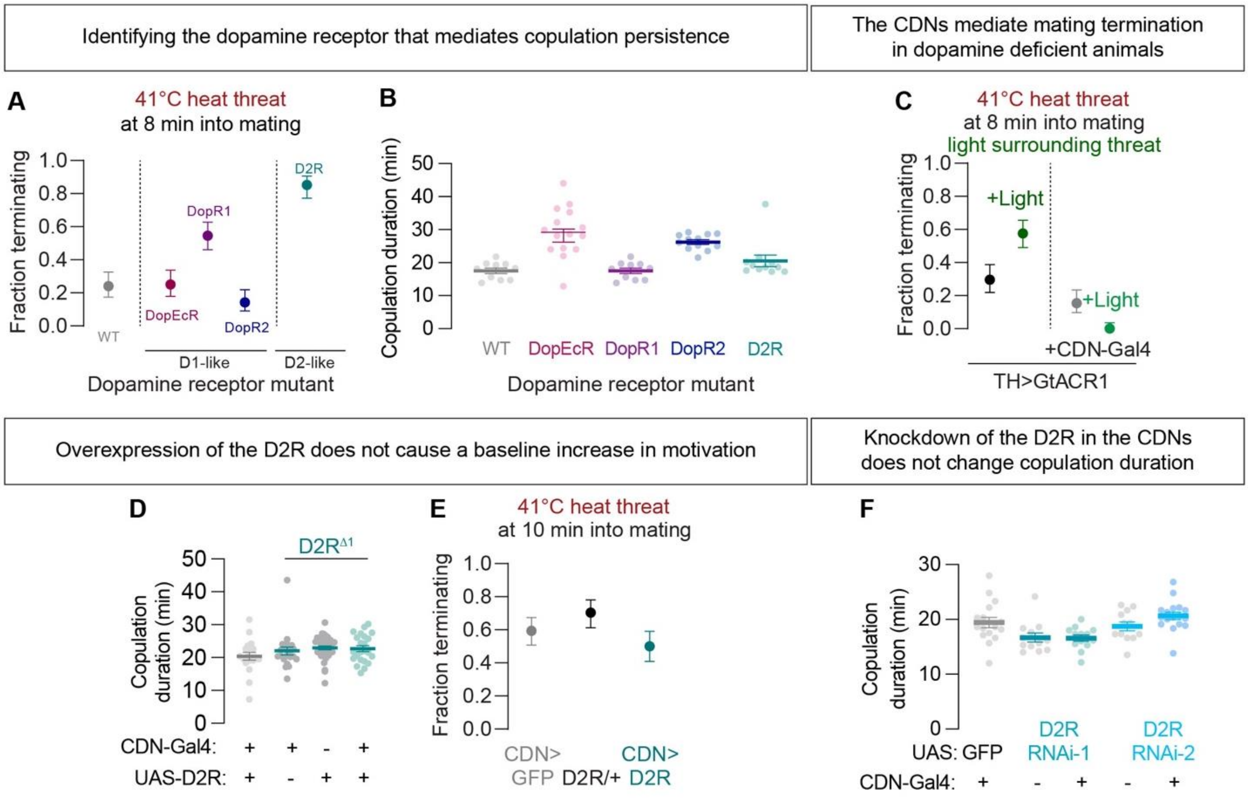

**Figure S4: Dopamine signaling to the CDNs through the D2R supports mating persistence**

**A.** Of the four dopamine receptors in Drosophila, whole-animal deletion of the D2 receptor most strongly increases termination in response to heat threats (n=27-33). Note: we observed a courtship deficit and low mating rate in D2R mutant flies when conducting these experiments not previously observed[6,17].

**B.** Mutation of any of the four known dopamine receptors in Drosophila males has little effect on copulation duration (n=10-15).

**C.** Electrical silencing of the dopamine neurons increases termination in response to a heat threat, an effect that is blocked by CDN silencing (n=26-33).

**D.** Deletion and rescue of D2R expression in the CDNs does not alter copulation duration (n=19-36).

**E.** Expression of UAS-D2R in the CDNs does not decrease the fraction of mating pairs terminating in response to a heat threat delivered at 10 minutes (n=27-32).

**F.** RNAi-mediated knockdown of D2R in the CDNs does not alter copulation duration (n=12-17).

#### Supplementary Figure 5

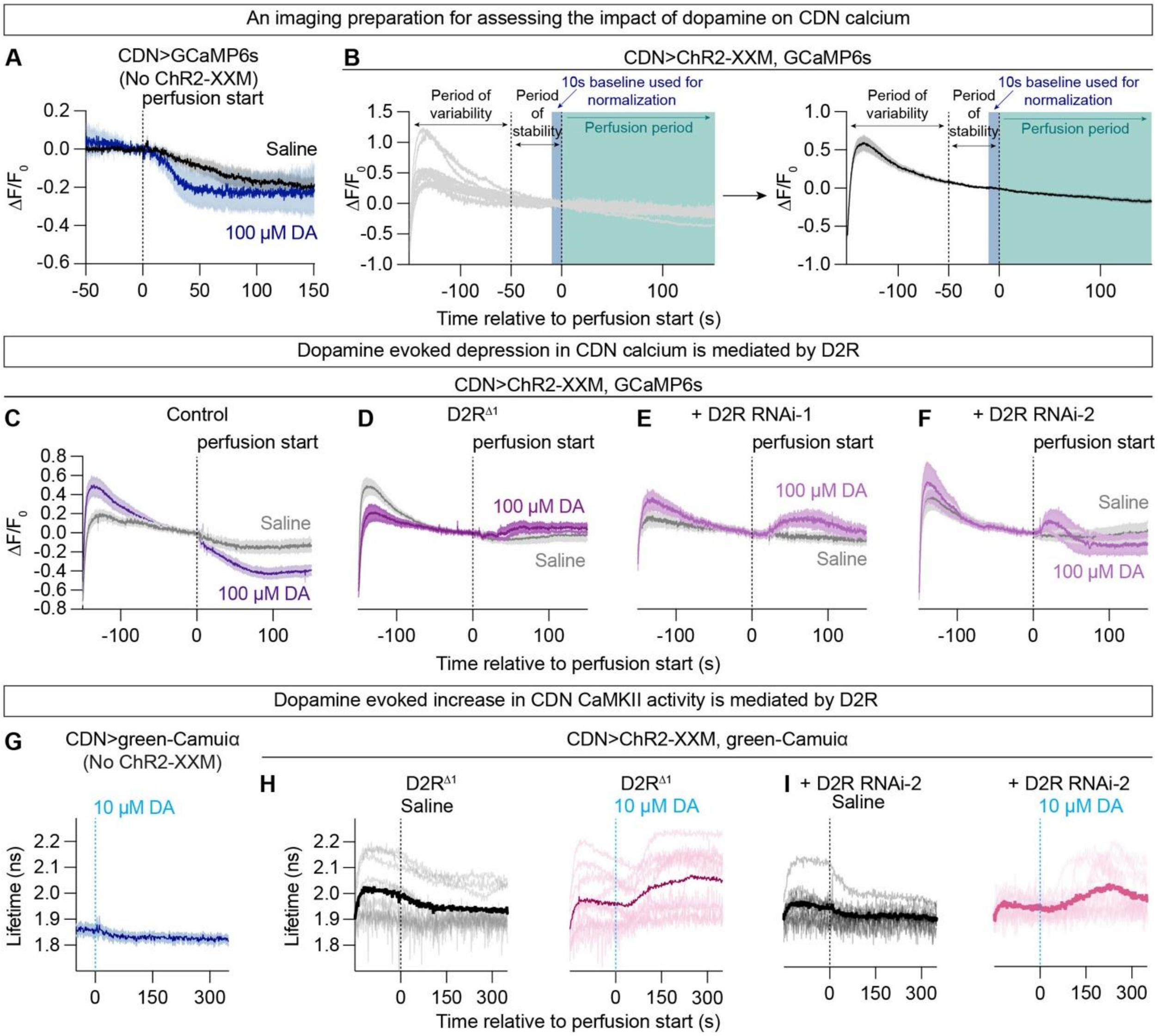

**Figure S5: Physiological evidence for D2R mediating the action of dopamine on the CDNs**

**A.** In the absence of ChR2-XXM driven stimulation, the effects of dopamine on CDN calcium are less obvious (n=10 each).

**B.** Example experiment where ChR2-XXM is used to elevate CDN activity before bath applying dopamine (left: individual traces, right: average trace displayed as mean ± SEM). Because the initial rise in calcium is variable, we wait until calcium levels stabilize before perfusing and use the 10 s immediately preceding the start of perfusion as a baseline. Traces in the main figures have been cropped to only show activity once calcium levels have stabilized. Control animals not provided any perfusion are shown (n=10 each).

**C.** Full trace of **Figure 2F**, left panel: CDN calcium signal is depressed upon bath application of 100 μM dopamine (n=10 each).

**D.** Whole animal D2R mutants do not show a depression in calcium signaling following bath application of 100 μM dopamine (n=10 each).

**E.** Full trace of **Figure 2F**, middle panel: animals with the D2R knockdown in the CDNs do not show depression of CDN calcium levels following bath application of 100 μM dopamine (n=10 each).

**F.** Animals with the D2R knocked down with a second, independent RNAi in the CDNs do not show depression of CDN calcium levels following bath application of 100 μM dopamine (n=10).

**G.** In the absence of ChR2-XXM driven stimulation dopamine application does not alter CaMKII activity in the CDNs (n=10 each).

**H.** D2R mutants show a weak increase in CaMKII activity following bath application of 10 μM dopamine (n=10 each).

**I.** Animals with the D2R knocked down with a second, independent RNAi in the CDNs show a weak increase in CaMKII activity following bath application of 10 μM dopamine (n=10 each).

#### Supplementary Figure 6

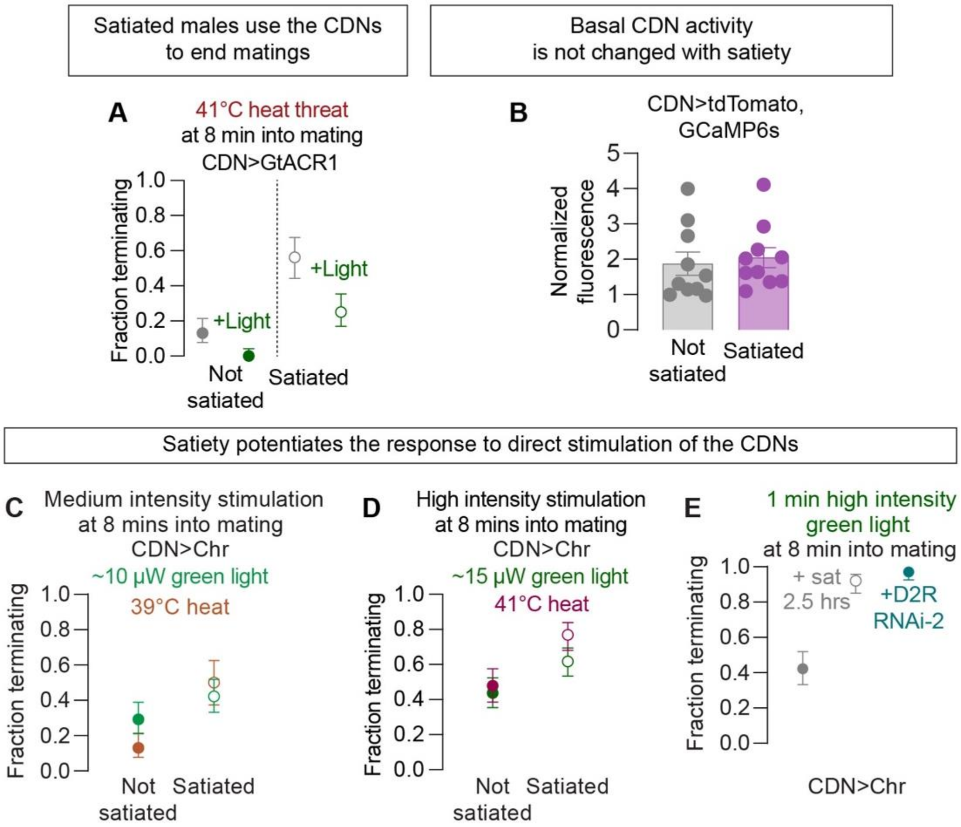

**Figure S6: Satiety sensitizes matings to CDN stimulation**

- **A.** Electrical silencing of the CDNs counteracts the satiety-induced increase in mating termination in response to a heat threat at 8 mins (n=16-23).
- **B.** Basal activity of the CDNs does not change with satiety state, as reported by normalizing GCaMP6s to tdTomato fluorescence (n=10 each).
- **C.** Satiety potentiates the response to a moderate heat threat similarly to moderate CDN activation (n=22-26).
- **D.** Satiety similarly potentiates the response to a strong heat threat and strong CDN activation (n=25-34).
- **E.** Satiety and D2R knockdown in the CDNs potentiate the response to direct stimulation of the CDNs similarly (n=25-33).

#### Supplementary Figure 7

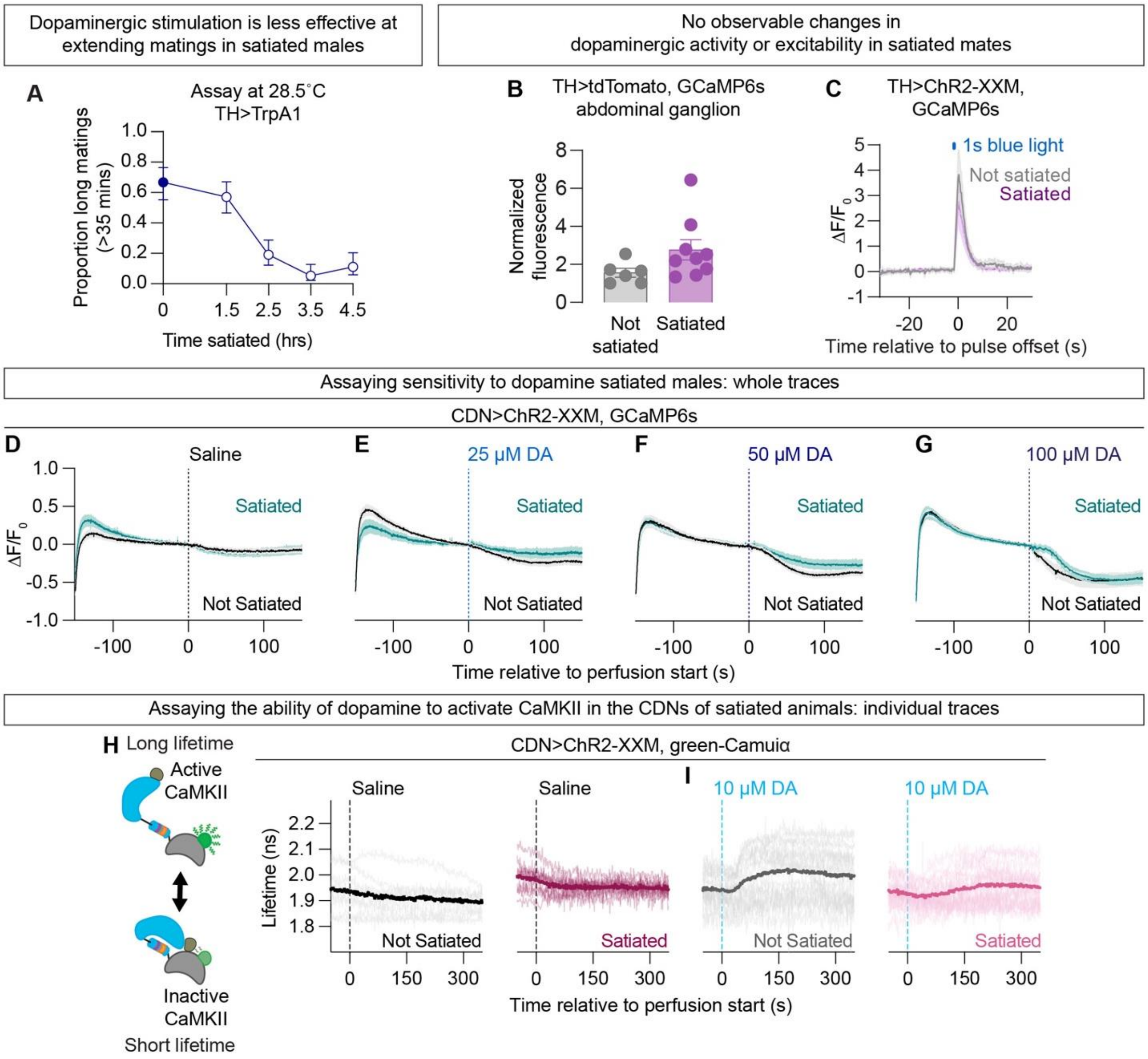

**Figure S7: Assessment of dopaminergic signaling in satiated animals**

- **A.** Dopaminergic stimulation becomes less effective at extending matings after males spend ~2.5 hours mating with females in the satiety assay (n=18-21). Note: the 2.5 hour time point is reproduced from **Figure 3A**.
- **B.** Basal activity of dopamine neurons in the abdominal ganglion does not change with satiety state, as reported by normalizing GCaMP6s to tdTomato fluorescence (n=6-9).
- **C.** Optogenetically evoked calcium activity in abdominal ganglion dopamine neurons does not change with satiety state (n=4-6).
- **D.** Full trace of **Figure 3B**, left panel: CDN calcium signal does not differ between not satiated and satiated animals (n=10 each).
- **E.** Full trace of **Figure 3B**, center left panel: CDN calcium signal is less depressed by bath application of 25 μM dopamine in satiated animals (n=10 each).
- **F.** Full trace of **Figure 3B**, center right panel: CDN calcium signal is less depressed by bath application of 50 μM dopamine in satiated animals (n=10 each).
- **G.** CDN calcium signal is depressed upon bath application of 100 μM dopamine in not satiated and satiated animals (n=10 each).

**H.** Left: schematic of the CaMKII sensor green-Camuiα. Right: Individual traces corresponding to **Figure 3D**, left panel: saline application does not increase CaMKII activity in not satiated or satiated flies (n=10 each).

**I.** Individual traces corresponding to **Figure 3D**, middle panel: bath application of 10 μM dopamine increases CaMKII activity more strongly in not satiated than satiated flies (n=10 each).

#### Supplementary Figure 8

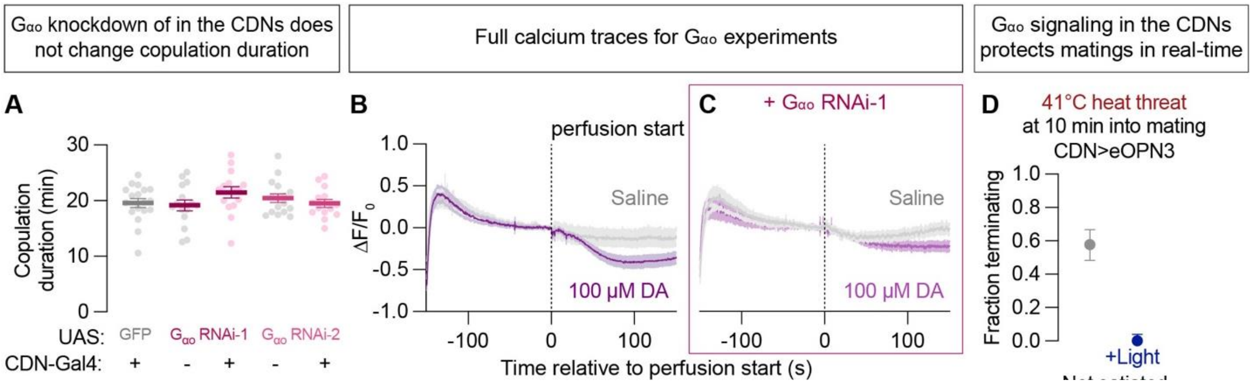

**Figure S8: Activation of $G_{\alpha o}$ signaling in the CDNs protects matings in real-time**

**A.** Knockdown of $G_{\alpha o}$ in the CDNs does not change copulation duration (n=8-18).

**B.** Full trace of Figure 4C, left panel, CDN calcium signal is depressed upon bath application of 100 µM dopamine in control animals (n=10 each).

**C.** Full trace of Figure 4C, middle panel, CDN calcium signal is only slightly depressed upon bath application of 100 µM dopamine in animals with $G_{\alpha o}$ knocked down in the CDNs (n=10 each).

**D.** Optogenetic activation of eOPN3 in the CDNs suppresses mating termination in response to a heat threat (n=24-26).

#### Supplementary Figure 9

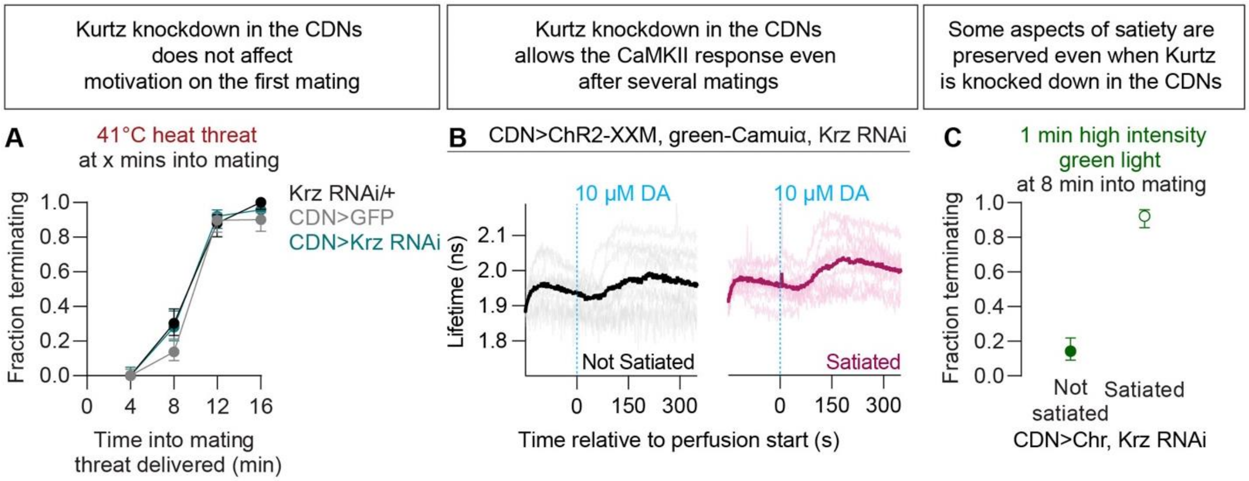

**Figure S9: Knockdown of β-arrestin in the CDNs prevents most aspects of satiety without altering baseline motivation**

**A.** Knockdown of β-arrestin (Kurtz) does not change motivational dynamics within a single mating (n=20-33).

**B.** Individual traces corresponding to **Figure 5C**: dopamine application activates CaMKII in not satiated and satiated animals when Kurtz is knocked down in the CDNs (n=10 each).

**C.** Knockdown of Kurtz does not block the satiety induced potentiation of the termination response to direct activation of the CDNs (n=25-26).

#### Supplementary Figure 10

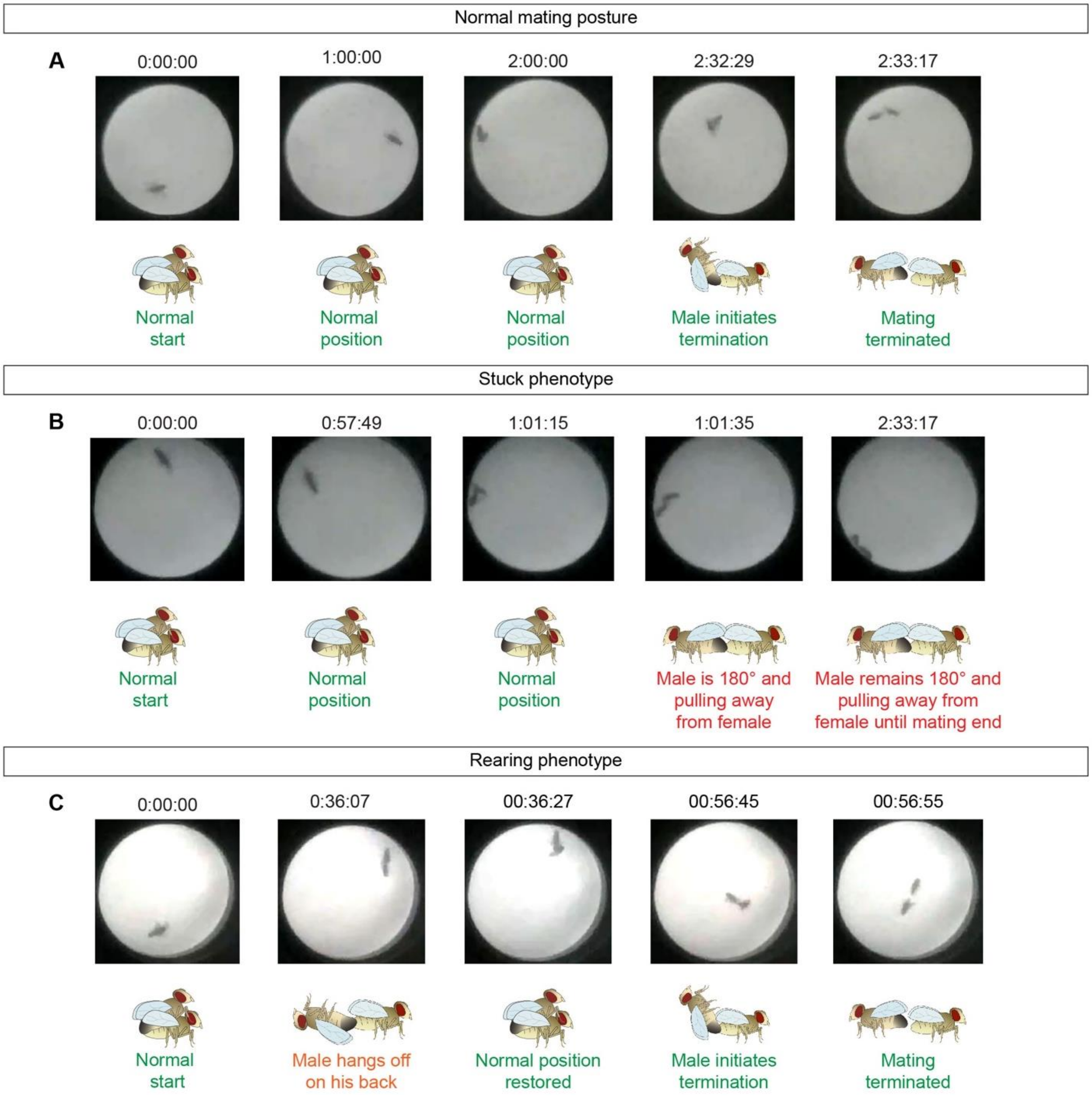

**Figure S10: Scoring extended copulations**

**A.** Depiction of flies maintaining normal mating posture throughout a long copulation until natural termination.

**B.** Depiction of flies displaying stuck behavior, in which the male is physically pulling away from the female but unable to terminate the copulation.

**C.** Depiction of flies displaying rearing behavior, in which the male temporarily increases the angle between himself and the female without turning to pull away before returning to normal mating posture. Time spent rearing in mating is counted towards overall copulation duration.

### SUPPLEMENTARY TABLES

#### Supplementary Table S1: Statistical Tests

Fisher's exact test and Mann Whitney U-test test were performed using Prism 10. All tests are unpaired with a p-value of alpha = 0.05. All tests with multiple comparisons use post hoc Bonferroni corrections: (0.05/n), where n is the number of unique comparisons made. Statistical significance is indicated in red.

| **Figure, Statistical test used** | **Null hypothesis** | **P-value<br>(alpha = 0.05)** |
|---|---|---|
| **FIGURE 1** | | |
| B – Fisher's exact test, groups numbered as:<br>1 = not satiated<br>2 = satiated<br>3 = satiated + 1 day recovery<br>4 = satiated + 2 day recovery<br>5 = satiated + 3 day recovery | No difference between termination probabilities | *Corrected significance value:*<br>0.0125<br><br>1-2: 0.0005<br>1-3: 0.0074<br>1-4: 0.2925<br>1-5: 0.2843<br>2-3: 0.3585<br>2-4: 0.0061<br>2-5: 0.0106<br>3-4: 0.0790<br>3-5: 0.1197<br>4-5: >0.9999 |
| C – Fisher's exact test | No difference between termination probabilities | 0.0004 |
| D – Fisher's exact test | No difference between termination probabilities | TH>TrpA1: <0.0001<br>TH>TrpA1 + Tsh-G80: 0.2416 |
| E – Fisher's exact test, groups numbered as:<br>1 = not satiated<br>2 = satiated without light<br>3 = satiated with light | No difference between termination probabilities | *Corrected significance value:*<br>0.025<br><br>1-2: 0.0005<br>1-3: >0.9999<br>2-3: 0.0021 |
| **FIGURE 2** | | |
| A – Fisher's exact test, groups numbered as:<br>1 = no light, +retinal<br>2 = no retinal, +light<br>3 = +retinal, +light | No difference between termination probabilities | *Corrected significance value:*<br>0.025<br><br>1-2: 0.0402<br>1-3: 0.0108<br>2-3: <0.0001 |
| C – Fisher's exact test, groups numbered as:<br>1 = CDN>UAS-D2R<br>2 = $D2R^{\Delta 1}$; UAS-D2R<br>3 = $D2R^{\Delta 1}$; CDN-Gal4<br>4 = $D2R^{\Delta 1}$; CDN>UAS-D2R | No difference between termination probabilities | *Corrected significance value:*<br>0.017<br><br>1-2: <0.0001<br>1-3: 0.0011<br>1-4: 0.5725<br>2-3: 0.3520<br>2-4: 0.0002<br>3-4: 0.0079 |

| | | |
|---|---|---|
| D – Fisher's exact test,<br>groups numbered as:<br>1 = CDN>myrGFP<br>2 = D2R RNAi/+<br>3 = CDN>D2R-RNAi | No difference between termination probabilities within each RNAi line | *Corrected significance value:*<br>0.025<br><br>*D2R-RNAi-1 (BDSC# 36824):*<br>1-3: <0.0001<br>2-3: <0.0001<br><br>*D2R-RNAi-2 (VDRC# 1820):*<br>1-3: <0.0001<br>2-3: 0.0027 |
| F – Mann Whitney U-test on the difference between before (-10 to 0s) and after (75 to 100s) perfusion | No difference between saline and dopamine perfusion values | No RNAi: 0.0007<br>D2R-RNAi-1: 0.0630 |
| H – Mann Whitney U-test on the difference between before (-10 to 0s) and after (150 to 300s) perfusion | No difference between saline and dopamine perfusion values | No RNAi: 0.0003<br>D2R-RNAi-1: 0.1045 |
| **FIGURE 3** | | |
| A – Fisher's exact test | No difference between the proportion of flies mating longer than 35 mins | <0.0001 |
| B – Mann Whitney U-test on the difference between before (-10 to 0s) and after (75 to 100s) perfusion | No difference between not satiated and satiated values | Saline: 0.5687<br>25 μM dopamine: 0.0603<br>50 μM dopamine: 0.0155 |
| C – Mann Whitney U-test on the difference between before (-10 to 0s) and after (150 to 300s) perfusion | No difference between not satiated and satiated values | Saline: 0.8286<br>10 μM dopamine: 0.0653 |
| **FIGURE 4** | | |
| B – Fisher's exact test,<br>groups numbered as:<br>1 = CDN>CD8-GFP<br>2 = Gαo RNAi/+<br>3 = CDN> Gαo-RNAi | No difference between termination probabilities within each RNAi line | *Corrected significance value:*<br>0.025<br><br>Gαo-*RNAi-1 (BDSC# 28010):*<br>1-3: <0.0001<br>2-3: <0.0001<br><br>Gαo-*RNAi-2 (BDSC# 34653):*<br>1-3: <0.0001<br>2-3: <0.0001 |
| C – Mann Whitney U-test on the difference between before (-10 to 0s) and after (75 to 125s) perfusion | No difference between saline and dopamine perfusion values | No RNAi: 0.0185<br>Gαo-RNAi-1: 0.1051 |
| E – Fisher's exact test | No difference between termination probabilities | 0.1874 |
| F – Mann Whitney U-test on the difference between baseline (0 to 2s) and after (75 to 100s) perfusion | No difference between values | Not satiated: 0.3930<br>Satiated: 0.2475 |
| G – Fisher's exact test | No difference between termination probabilities | 0.2326 |

| | | |
|---|---|---|
| H – Fisher's Exact Test | No difference between termination probabilities | 0.2326 |
| **FIGURE 5** | | |
| B – Fisher's exact test | No difference between values | CDN>GFP: 0.0179<br>Krz RNAi/+: 0.0204<br>CDN>Krz RNAi: >0.9999 |
| C – Mann Whitney U-test on the difference between before (-10 to 0s) and after (150 to 300s) perfusion | No difference between values | 0.5787 |
| **SUPPLEMENTARY FIGURE 1** | | |
| A – Mann Whitney U-test | No difference between initial and end percentage of mating behaviors | 0.0079 |
| B – Fisher's exact test, groups numbered as:<br>1 = not satiated<br>2 = satiated 1.5 hrs<br>3 = satiated 2.5 hrs<br>4 = satiated 3.5 hrs<br>5 = satiated 4.5 hrs | No difference between termination probabilities | *Corrected significance value:*<br>0.0125<br><br>1-2: 0.0165<br>1-3: <0.0001<br>1-4: <0.0001<br>1-5: <0.0001 |
| C – Mann-Whitney U-test, groups numbered as:<br>1 = not satiated<br>2 = satiated 2.5 hrs<br>3 = satiated 4.5 hrs | No difference between copulation durations | *Corrected significance value:*<br>0.025<br><br>1-2: 0.2563<br>1-3: 0.4803<br>2-3: 0.6439 |
| D – Fisher's exact test, groups numbered as:<br>1 = not satiated<br>2 = W1118<br>3 = Chr-mVenus<br>4 = Elav>SP | No difference between termination probabilities | *Corrected significance value:*<br>0.017<br><br>1-2: 0.0385<br>1-3: 0.0339<br>1-4: 0.4512 |
| F – Mann Whitney U-test | No difference between ejaculatory bulb fullness scores | <0.0001 |
| G – Fisher's Exact Test | No difference between termination probabilities | 0.6092 |
| H – Mann Whitney U-test, groups numbered as:<br>1 = not satiated<br>2 = satiated, no light<br>3 = satiated, light | No difference between ejaculatory bulb scores | *Corrected significance value:*<br>0.025<br><br>1-2: <0.0001<br>1-3: >0.9999<br>2-3: <0.0001 |
| I – Fisher's Exact Test, groups numbered as:<br>1 = not satiated<br>2 = satiated, no light<br>3 = satiated, light | No difference between termination probabilities | *Corrected significance value:*<br>0.025<br><br>1-2: 0.0012<br>1-3: 0.0056<br>2-3: 0.7892 |

| | | |
|---|---|---|
| **SUPPLEMENTARY FIGURE 2** | | |
| C – Fisher's Exact Test | No difference between termination probabilities within each genotype | TrpA1/+: 0.5008<br>P1>TrpA1: 0.6138<br>NPF>TrpA1: 0.2097<br>pCd>TrpA1: 0.1651<br>NPF, pCd>TrpA1: 0.0193<br>mAL>TrpA1: 0.6782<br>vAB3>TrpA1: 0.5798<br>PPN1>TrpA1: >0.9999<br>NP5945>TrpA1: 0.3391<br>VT02857>TrpA1: 0.0938<br>VT02857, NP5945>TrpA1: 0.4693 |
| D – Fisher's Exact Test | No difference between termination probabilities | CRN>TrpA1: <0.0001 |
| E – Mann Whitney U-test | No difference in copulation duration | CRN>TrpA1: 0.4066 |
| **SUPPLEMENTARY FIGURE 3** | | |
| A – Fisher's Exact Test | No difference between termination probabilities within each genotype | TH>GFP: 0.4902<br><br>TrpA1/+ 23°C data was also used in S2C<br>*Corrected significance value:*<br>0.025<br><br>TrpA1/+: >0.9999 |
| C – Mann Whitney U-test | No difference in copulation duration within each genotype | TH>GFP: 0.0322<br>TrpA1/+: 0.0032<br>TH>TrpA1: 0.7226<br>TH>TrpA1 + Tsh-Gal80: 0.0032 |
| D – Fisher's Exact Test | No difference between the proportion of flies mating longer than 35 mins within each genotype | TH>GFP: >0.9999<br>TrpA1/+: >0.9999<br>TH>TrpA1: <0.0001 |
| E – Fisher's Exact Test | No difference between termination probabilities | 0.0098 |
| F – Mann Whitney U-test | No difference between the number of matings | 0.8500 |
| G – Mann Whitney U-test, groups numbered as:<br>1 = no light<br>2 = no retinal<br>3 = light and retinal | No difference between copulation durations | *Corrected significance value:*<br>0.025<br><br>1-2: 0.0029<br>1-3: 0.9482 |
| H – Fisher's Exact Test | No difference between termination probabilities | >0.9999 |
| **SUPPLEMENTARY FIGURE 4** | | |
| A – Fisher's Exact Test, groups numbered as:<br>1 = WT<br>2 = DopEcR<br>3 = DopR1 | No difference between termination probabilities | *Corrected significance value:*<br>0.0125<br><br>1-2: 0.1430<br>1-3: <0.0001 |

| | | |
|---|---|---|
| 4 = DopR2<br>5 = D2R | | 1-4: 0.6695<br>1-5: <0.0001 |
| B – Mann Whitney U-test, groups numbered as:<br>1 = WT<br>2 = DopEcR<br>3 = DopR1<br>4 = DopR2<br>5 = D2R$^{\Delta 1}$ | No difference between copulation durations | *Corrected significance value:*<br>0.0125<br><br>1-2: <0.0001<br>1-3: >0.9999<br>1-4: <0.0001<br>1-5: 0.2890 |
| C – Fisher's Exact Test, groups numbered as:<br>1 = TH>ACR, no light<br>2 = TH>ACR, light<br>3 = TH, CDN>ACR, no light<br>4 = TH, CDN>ACR, light | No difference between termination probabilities | *Corrected significance value:*<br>0.025<br><br>1-2: 0.0392<br>1-3: 0.3265<br>2-4: <0.0001<br>3-4: 0.1104 |
| D – Mann Whitney U-test, groups numbered as:<br>1 = CDN>UAS-D2R<br>2 = D2R$^{\Delta 1}$, CDN/+<br>3 = D2R$^{\Delta 1}$, UAS-D2R/+<br>4 = D2R$^{\Delta 1}$, CDN>UAS-D2R | No difference between copulation durations | *Corrected significance value:*<br>0.017<br><br>1-2: 0.3334<br>1-3: 0.0106<br>1-4: 0.1721<br>2-4: 0.3701<br>3-4: 0.6855 |
| E – Fisher's Exact Test, groups numbered as:<br>1 = CDN>GFP<br>2 = UAS-D2R/+<br>3 = CDN>UAS-D2R | No difference between termination probabilities | *Corrected significance value:*<br>0.025<br><br>1-3: 0.6039<br>2-3: 0.1707 |
| F – Mann Whitney U-test, groups numbered as:<br>1 = CDN>GFP<br>2 = D2R-RNAi/+<br>3 = CDN>D2R-RNAi | No difference between copulation durations | *Corrected significance value:*<br>0.025<br><br>*D2R-RNAi-1 (BDSC# 36824):*<br>1-3: 0.0127<br>2-3: 0.6781<br><br>*D2R-RNAi-2 (VDRC# 1820):*<br>1-3: 0.1115<br>2-3: 0.1247 |
| **SUPPLEMENTARY FIGURE 5** | | |
| A – Mann Whitney U-test on the difference between before (-10 to 0s) and after (75 to 100s) perfusion | No difference between saline and dopamine perfusion values | 0.7394 |
| C – Mann Whitney U-test on the difference between before (-10 to 0s) and after (75 to 100s) perfusion | No difference between saline and dopamine perfusion values | Same statistics for Figure 2F:<br>No RNAi: 0.0007 |
| D – Mann Whitney U-test on the difference between before (-10 to 0s) and after (75 to 100s) perfusion | No difference between saline and dopamine perfusion values | 0.0753 |
| E – Mann Whitney U-test on the difference between before (-10 to 0s) and after (75 to 100s) perfusion | No difference between saline and dopamine perfusion values | Same statistics for Figure 2F:<br>D2R-RNAi-1: 0.0630 |

| | | |
|---|---|---|
| F – Mann Whitney U-test on the difference between before (-10 to 0s) and after (75 to 100s) perfusion | No difference between saline and dopamine perfusion values | 0.3527 |
| H – Mann Whitney U-test on the difference between before (-10 to 0s) and after (150 to 300s) perfusion | No difference between saline and dopamine perfusion values | 0.0005 |
| I – Mann Whitney U-test on the difference between before (-10 to 0s) and after (150 to 300s) perfusion | No difference between saline and dopamine perfusion values | 0.1045 |
| **SUPPLEMENTARY FIGURE 6** | | |
| A – Fisher's Exact Test, groups numbered as:<br>1 = not satiated, no light<br>2 = not satiated, light<br>3 = satiated, no light<br>4 = satiated, light | No difference between termination probabilities | *Corrected significance value:*<br>0.017<br><br>1-2: 0.4884<br>1-3: 0.0029<br>1-4: 0.2289<br>2-4: 0.0182<br>3-4: 0.0874 |
| B – Mann Whitney U-test | No difference between normalized fluorescence values | 0.3527 |
| C – Fisher's Exact Test, groups numbered as:<br>1 = not satiated, 10 μW green light<br>2 = satiated, 10 μW green light<br>3 = not satiated, 39°C heat threat<br>4 = satiated, 39°C heat threat | No difference between termination probabilities | *Corrected significance value:*<br>0.025<br><br>1-2: 0.3876<br>1-3: 0.2865<br>3-4: 0.0230<br>2-4: >0.9999 |
| D – Fisher's Exact Test, groups numbered as:<br>1 = not satiated, 15 μW green light<br>2 = satiated, 15 μW green light<br>3 = not satiated, 41°C heat threat<br>4 = satiated, 41°C heat threat | No difference between termination probabilities | *Corrected significance value:*<br>0.025<br><br>1-2: 0.2171<br>1-3: 0.7937<br>3-4: 0.0448<br>2-4: 0.2684 |
| E – Fisher's Exact Test, groups numbered as:<br>1 = CDN>Chr, not satiated<br>2 = CDN>Chr, satiated<br>3 = CDN>Chr, D2R-RNAi-2, not satiated | No difference between termination probabilities | *Corrected significance value:*<br>0.025<br><br>1-2: 0.0002<br>1-3: <0.0001<br>2-3: 0.5722 |
| **SUPPLEMENTARY FIGURE 7** | | |
| A – Fisher's Exact Test, groups numbered as:<br>1 = not satiated<br>2 = satiated 1.5 hrs<br>3 = satiated 2.5 hrs<br>4 = satiated 3.5 hrs<br>5 = satiated 4.5 hrs | No difference between the proportion of flies mating longer than 35 mins within each genotype | *Corrected significance value:*<br>0.0125<br><br>1-2: 0.7424<br>1-3: 0.0038<br>1-4: 0.0001<br>1-5: 0.0016 |
| B – Mann Whitney U-test | No difference between normalized fluorescence values | 0.0663 |

| | | |
|---|---|---|
| C – Mann Whitney U-test | No difference between not satiated and satiated values | peak: 0.3524<br>residual (15-20s after pulse): 0.4762 |
| D – Mann Whitney U-test on the difference between before (-10 to 0s) and after (75 to 100s) perfusion | No difference between not satiated and satiated values | Same statistics as Figure 3B:<br>Saline: 0.5687 |
| E – Mann Whitney U-test on the difference between before (-10 to 0s) and after (75 to 100s) perfusion | No difference between not satiated and satiated values | Same statistics as Figure 3B:<br>25 µM dopamine: 0.0603 |
| F – Mann Whitney U-test on the difference between before (-10 to 0s) and after (75 to 100s) perfusion | No difference between not satiated and satiated values | Same statistics as Figure 3B:<br>50 µM dopamine: 0.0155 |
| G – Mann Whitney U-test on the difference between before (-10 to 0s) and after (75 to 100s) perfusion | No difference between not satiated and satiated values | 0.5787 |
| H – Mann Whitney U-test on the difference between before (-10 to 0s) and after (150 to 300s) perfusion | No difference between not satiated and satiated lifetime values | Same statistics as Figure 3C:<br>Saline: 0.8286 |
| I – Mann Whitney U-test on the difference between before (-10 to 0s) and after (150 to 300s) perfusion | No difference between not satiated and satiated lifetime values | Same statistics as Figure 3C:<br>10 µM dopamine: 0.0653 |
| **SUPPLEMENTARY FIGURE 8** | | |
| A – Mann Whitney U-test, groups numbered as:<br>1 = CDN>GFP<br>2 = Gαo-RNAi<br>3 = CDN> Gαo-RNAi | No difference between copulation durations | *Corrected significance value:*<br>0.025<br><br>Gαo-*RNAi-1 (BDSC# 28010):*<br>1-3: 0.0877<br>2-3: 0.1286<br><br>Gαo-*RNAi-2 (BDSC# 34653):*<br>1-3: 0.5674<br>2-3: 0.4916 |
| B – Mann Whitney U-test on the difference between before (-10 to 0s) and after (75 to 125s) perfusion | No difference between saline and dopamine perfusion values | Same statistics as Figure 4C:<br>No RNAi: 0.0185 |
| C – Mann Whitney U-test on the difference between before (-10 to 0s) and after (75 to 125s) perfusion | No difference between saline and dopamine perfusion values | Same statistics as Figure 4C:<br>Gαo-RNAi-1: 0.1051 |
| D – Fisher's Exact Test | No difference between termination probabilities | <0.0001 |
| **SUPPLEMENTARY FIGURE 9** | | |
| A – Fisher's Exact Test, groups numbered as:<br>1 = CDN>GFP<br>2 = Krz RNAi/+<br>3 = CDN>Krz RNAi | No difference between termination probabilities within a time into mating | *Corrected significance value:*<br>0.025<br><br>*4 mins into mating:*<br>1-3: >0.9999<br>2-3: >0.9999<br><br>*8 mins into mating:*<br>1-3: 0.3104<br>2-3: >0.9999 |

| | | |
|---|---|---|
| | | *12 mins into mating:*<br>1-3: >0.9999<br>2-3: >0.9999<br><br>*12 mins into mating:*<br>1-3: 0.6204<br>2-3: >0.9999 |
| B – Mann Whitney U-test on the difference between before (-10 to 0s) and after (150 to 300s) perfusion | No difference between not satiated and satiated lifetime values | Same statistics as Figure 5C: 0.5787 |
| C – Fisher's Exact Test | No difference between termination probabilities | <0.0001 |

#### Supplementary Table S2: Antibodies

| Antibody | Source | Identifier | Concentration | Figure(s) |
|---|---|---|---|---|
| Mouse anti-TH | Immunostar | 22941 | 1:500 | 2B |
| Chicken anti-GFP | Aves Labs | GFP-1010 | 1:1000 | 2B, S3B |
| Mouse anti-brunchpilot (nc82) | Developmental Studies Hybridoma Bank | N/A | 1:7 | S3B |
| Donkey anti-mouse 647 | Jackson ImmunoResearch | 711-605-151 | 1:200, 1:100 | 2B, S3B |
| Donkey anti-chicken 488 | Jackson ImmunoResearch | 703-545-155 | 1:1000, 1:200 | 2B, S3B |

#### Supplementary Table S3: Chemicals

| Reagent | Source | Identifier |
|---|---|---|
| All-*trans*-retinal | Sigma Aldrich | R2500 |
| All-*trans*-retinal | Spectrum Chemical | R3041 |
| Dopamine hydrochloride | Tocris | 3548 |
| Sodium chloride | Fisher Chemical | S271-500 |
| Potassium chloride | Alpha Aesar | A11661.0B |
| TES | Thermo Fisher Scientific | B21819.18 |
| Trehalose, Dihydrate | EMD Millpore Corp. | 625625-50GM |
| (D)-+-Glucose | Sigma Aldrich | G8270-1KG |
| Sodium bicarbonate | Acors Organics | 447102500 |
| Sodium phosphate | Research Products International | S23120-500.0 |
| Magnesium chloride | Fisher Bioreagents | BP214-500 |
| Calcium chloride | Fisher Chemical | C89-500 |
| Phosphate Buffered Saline 10X Molecular Biology Grade | Mediatech, Inc. | 46-013-CM |
| Tritron X-100 Scintillation Grade | Research Products International | 111036 |
| Potassium Chloride | Molecular Toxicology | 26-645 |
| Normal Donkey Serum | Jackson ImmunoResearch | AB_2337258 |
| Paraformaldehyde 16% | Polysciences | 18814 |

#### Supplementary Table S4: Other materials

| Resource | Source | Identifier |
|---|---|---|
| **Miscellaneous** | | |
| Petri Dish 35 x 10 mm | VWR | 10799-192 |
| Formula 4-24 Instant Drosophila Medium, Blue | Carolina Biological Supply | 173210 |
| 1.5 mL black Eppendorf tubes | Fisher Scientific | 15386548 |
| 50 mL Falcon tubes | Corning | 352098 |
| Dissection microscope | Nikon | SMZ745T |
| Excelis HDMI camera | ACCU-SCOPE | AU-600-HD |
| Glass slides | Azer Scientific | EMS200WS |
| Cover slips | VWR | 48393-026 |
| Cover slips (used to make a bridge for mounting) | VWR | 16004-094 |
| Vacuum grease (used to make a bridge for mounting) | Dow Corning | 2021854-0807 |
| Clear nail polish (to seal slides) | Sally Hansen | N/A |
| ProLong Diamond Antifade Mountant | ThermoFisher Invitrogen | P36961 |
| **Links to resources for behavioral arenas** | | |
| Behavior arenas for termination assays | Thornquist et. al, 2020; updated for this paper | https://github.com/CrickmoreRoguljaLabs/Waterworks |
| Copulation duration 32-well plates | Boutros, Miner et. al, 2017 | https://github.com/CrickmoreRoguljaLabs/Fly-mating-behavior-arena-designs |
| Green Light Pads | This publication | https://github.com/CrickmoreRoguljaLabs/GreenLightpads |
| Thermogenetics Box | This publication | https://github.com/CrickmoreRoguljaLabs/ThermogeneticsBox |

#### Supplementary Table S5: Software

| Software | Source | Identifier |
|---|---|---|
| MATLAB Jeffery's confidence interval code | Zhang et. al, 2018; updated for this paper | https://github.com/CrickmoreRoguljaLabs/ UpdatedJeffi |
| MATLAB copulation duration code | Thornquist et. al, 2020 | https://github.com/CrickmoreRoguljaLabs/FlyKnight |
| FLIMage software | Laviv et. al, 2020 | https://github.com/ryoheiyasuda/FLIMage_public |
| MATLAB calcium imaging anaylsis code | Gautham et. al, in revision | https://github.com/CrickmoreRoguljaLabs/GCaMP-from-FLIMage |
| Python FLIM analysis code | Gautham et. al, in revision | https://github.com/CrickmoreRoguljaLabs/flim-analysis |
| MATLAB GCaMP normalized to tdTomato code | Zhang et. al, 2016; updated for this paper | https://github.com/CrickmoreRoguljaLabs/GCaMP-tdt_Analysis |

#### Supplementary Table S6: Organisms/Strains

*All strains are D. melanogaster*

| Strain | Detailed Strain | Source | Identifier |
|---|---|---|---|
| W1118 with hs-hid on the Y | $w^{1118}$/Dp(2;Y), P{hs-hid}Y | Bloomington Drosophila Stock Center | BDSC 24638 |
| UAS-CsChrimson-mVenus with hs-hid on the Y | $w^{1118}$, P{ 20XUAS-IVS-CsChrimson.mVenus}attp18/Dp(2;Y), P{hs-hid}Y | Bloomington Stock Center/Michael Crickmore Lab | BDSC 55134 (without hs-hid) |
| Wild-type Canton S | | Keleman et. al. 2012 | Flybase FBsn0000274 |
| NP2719-Gal4 | w*;P{GawB}GC17646$^{NP2719}$/CyO | *Drosophila* Genome Resource Center | DGRC 113024 |
| RepoGal80 | | Tzumin Lee Lab | N/A |
| TH-Gal4 | w[*];; P{ple-GAL4.F}3 | Bloomington Stock Center | BDSC 8848 |
| UAS-TrpA1 | w*; P{UAS-TrpA1(B).K}attP16 | Bloomington Stock Center | BDSC 26263 |
| tshGal80 | $w^{1118}$; P{GAL80}tsh$^{GAL80}$ | Julie Simpson Lab | N/A |
| UAS-GtACR1-eYFP (III) | $w^{1118}$;; P{UAS-IVS-GtACR1}attP2 | Adam Claridge/Chang Lab | BDSC 92983 |
| D2R$^{\Delta1}$ | Df(1)Dop2R[Delta1],Dop2R[Delta1] | Bloomington Stock Center | BDSC 78795 |
| D2R$^{\Delta1}$ with UAS-Dop2R.S | Df(1)Dop2R[Delta1],Dop2R[Delta1];; P{ UAS-Dop2R.S}3 | Bloomington Stock Center | BDSC 78797 |
| UAS-Dop2R.S | $w^{1118}$;; P{ UAS-Dop2R.S}3 | Bloomington Stock Center/Michael Crickmore Lab | Isolated from BDSC 78798 |
| UAS-myrGFP | $w^{1118}$;; P{UAS-IVS-myr::GFP}attP2 | Bloomington Stock Center | BDSC 32197 |
| UAS-CD8-GFP | $w^{1118}$;; P{UAS-mCD8::GFP.L}attP2 | Lee and Luo, 1999 | |
| UAS-Dicer2 | $w^{1118}$, P{UAS-Dcr-2.D}1 | Bloomington Stock Center | BDSC 24646 |
| UAS-D2R-RNAi-1 | $y^1sc^*v^1sev^{21}$; P{TRiP.GL01057}attP2 | Bloomington Stock Center | BSDC 36824 |
| UAS-D2R-RNAi-2 | $w^{1118}$; P{GD733}v1820 | Vienna Drosophila Resource Center | VDRC v1820 |
| UAS-ChR2-XXM | PBac{UAS-ChR2.XXM} | Robert Kittel Lab | N/A |
| UAS-GCaMP6s (III) | w*;;UAS-GCaMP6s | David Anderson Lab | N/A |
| UAS-green-camuiα | w*;;UAS-green-camuiα | Michael Crickmore Lab | N/A |
| UAS-$G_{\alpha o}$-RNAi-1 | $y^1v^1$; P{TRiP.JF02844}attP2 | Bloomington Stock Center | BDSC 28010 |
| UAS-$G_{\alpha o}$-RNAi-2 | $y^1sc^*v^1sev^{21}$; P{TRiP.HMS01129}attP2 | Bloomington Stock Center | BDSC 34653 |
| UAS-eOPN3 | yw*;; P{20XUAS-eOPN3.HA} | Stephen Thornquist/Michael Crickmore Lab | N/A |
| UAS-Krz-RNAi | $y^1v^1$; P{TRiP.HM05200}attP2 | Bloomington Drosophila Stock Center | BDSC 29523 |
| Elav-Gal4 | P{GawB}elav$^{c155}$ | Bloomington Stock Center | BDSC 458 |

| | | | |
|---|---|---|---|
| UAS-SP (sex peptide) | $w^{1118}$;; UAS-SP | Nakayama et. al, 1997 | |
| Crz-Gal4 | $w^{1118}$; Crz-Gal4 | Tayler et. al, 2012 | |
| P1-Gal4 | $w^{1118}$;; P{GMR71G01-GAL4}attP2 | Bloomington Stock Center | BDSC 39599 |
| NPF-Gal4 | $w^{1118}$, $y^{1}$ ;; P{NPF-Gal4.1}1 | Bloomington Stock Center | BDSC 25682 |
| pCd-Gal4 | $w^{1118}$;; P{GMR41A01-GAL4}attP2 | Bloomington Stock Center | BDSC 39425 |
| mAL-Gal4 | $w^{1118}$;; P{GMR43D01-GAL4}attP2 | Bloomington Stock Center | BDSC 64345 |
| vAB3-Gal4 | $y^{1}$ , w*;; P{GawB}Abd-BLDN/TM6B, $Tb^{1}$ | Bloomington Stock Center | BDSC 55848 |
| PPN1-Gal4 | $w^{1118}$;; P{GMR56C09-GAL4}attP2 | Bloomington Stock Center | BDSC 39145 |
| NP5945-Gal4 | y*, w*;; P{w[+mW.hs] = GawB}mura[NP5945]/TM6, P{w[-] = UAS-lacZ.UW23-1}UW23-1 | Kyoto Drosophila Genomics and Genetic Resources | DGGR 105062 |
| VT02857-Gal4 | $w^{1118}$;; P{VT002857-Gal4}attP2 | Vienna Drosophila Resource Center | VDRC 200494 |
| CRN-Gal4 | $w^{1118}$; P{GMR42G02-Gal4}attP40 | Zhou et. al, 2014 | N/A |
| DopEcR mutant | $w^{1118}$; PBac{PB}$DopEcR^{c02142}$ | Bloomington Stock Center | BDSC 10847 |
| DopR1 mutant | +;; $DopR1^{attp}$ | Barry Disckson Lab, Keleman et. al, 2012 | FBal0283278 |
| DopR2 mutant | +;; $DopR2^{attp}$ | Barry Disckson Lab, Keleman et. al, 2012 | FBal0283280 |
| $D2R^{P5}$ | $w^{1118}$ PBac{WH}$D2R^{f06521}$ | Bloomington Stock Center | BDSC 85250 |
| $D2R^{\Delta 2}$ | Df(1)Dop2R[Delta2],Dop2R[Delta2] | Bloomington Stock Center | BDSC 78796 |
| UAS-tdTomato | w*;; P{10XUAS-IVS-myr::tdTomato}attp2 | Bloomington Stock Center | BDSC 32221 |
| UAS-opGCaMP6s (II) | $w^{1118}$; P{20xUAS-IVS-Syn21-opGCaMP6s} attp40 | Bloomington Stock Center | BDSC 42746 |
| UAS-Chrimson | $w^{1118}$;; P{20xUAS-IVS-Syn21-CsChrimsontdTomato}attP2 | David Anderson Lab | N/A |

#### Supplementary Table S7: Genotypes and number of flies per experiment

*All strains are D. melanogaster*

*All females used in satiety assays are* $w^{1118}/w^{1118}$; +/+; +/+

*All females used as mating partners in behavioral experiment are* $w^{1118}$, UAS-Chrimson-mVenus; +/+; +/+ unless otherwise noted

*All males are unsatiated unless otherwise noted*

*All threats are 1 minute long*

| Figure | Label | Genotype (male) | Condition | N |
|---|---|---|---|---|
| **FIGURE 1** | | | | |
| A | - | - | - | - |
| B | Not satiated | Wildtype-Canton S | Not satiated, 41°C heat threat at 8 mins | 29 |
| | Satiated | Wildtype-Canton S | Satiated 2.5 hrs, 41°C heat threat at 8 mins | 21 |
| | Satiated + 1 day recovery | Wildtype-Canton S | Satiated 2.5 hrs + 1 day recovery, 41°C heat threat at 8 mins | 28 |
| | Satiated + 2 day recovery | Wildtype-Canton S | Satiated 2.5 hrs + 2 day recovery, 41°C heat threat at 8 mins | 35 |
| | Satiated + 3 day recovery | Wildtype-Canton S | Satiated 2.5 hrs + 3 day recovery, 41°C heat threat at 8 mins | 32 |
| C | Not satiated | $w^{1118}$/Y; UAS-TrpA1/+; +/+ | Not satiated, blue light stressor at 10 mins | 30 |
| | Satiated | $w^{1118}$/Y; UAS-TrpA1/+; +/+ | Satiated 4.5 hrs, blue light stressor at 10 mins | 30 |
| D | TH>TrpA1, Not stimulated | $w^{1118}$/Y; UAS-TrpA1/+; TH-Gal4/+ | Spent 4.5 hrs at 23°C prior to assay, blue light stressor at 10 mins | 27 |
| | TH>TrpA1, Pre-stimulated | $w^{1118}$/Y; UAS-TrpA1/+; TH-Gal4/+ | Spent 4.5 hrs at 28.5°C prior to assay, blue light stressor at 10 mins | 23 |
| | TH>TrpA1 + Tsh-Gal80, Not stimulated | $w^{1118}$/Y; UAS-TrpA1/Tsh-Gal80; TH-Gal4/+ | Spent 4.5 hrs at 23°C prior to assay, blue light stressor at 10 mins | 37 |
| | TH>TrpA1 + Tsh-Gal80, Pre-stimulated | $w^{1118}$/Y; UAS-TrpA1/Tsh-Gal80; TH-Gal4/+ | Spent 4.5 hrs at 28.5°C prior to assay, blue light stressor at 10 mins | 25 |
| E | TH>GtACR1, Not satiated | $w^{1118}$/Y; +/+; TH-Gal4/UAS-GtACR1 | Spent 2.5 hrs in dim light + 24 hrs | 30 |

| | | | | |
|---|---|---|---|---|
| | | | recovery, 41°C heat threat at 8 mins | |
| | TH>GtACR1, Satiated (grey) | $w^{1118}$/Y; +/+; TH-Gal4/UAS-GtACR1 | Satiated 2.5 hrs in dim light + 24 hrs recovery, 41°C heat threat at 8 mins | 15 |
| | TH>GtACR1, Satiated (green +Light) | $w^{1118}$/Y; +/+; TH-Gal4/UAS-GtACR1 | Satiated 2.5 hrs in dim light except when mating (then in green light) + 24 hrs recovery, 41°C heat threat at 8 mins | 25 |
| **FIGURE 2** | | | | |
| A | TH>GtACR1, no light | $w^{1118}$/Y; +/+; TH-Gal4/UAS-GtACR1 | Fed retinal, but no light during testing, 41°C heat threat at 8 mins | 31 |
| | TH>GtACR1, no retinal | $w^{1118}$/Y; +/+; TH-Gal4/UAS-GtACR1 | Not fed retinal, light surrounding threat, 41°C heat threat at 8 mins | 30 |
| | TH>GtACR1 | $w^{1118}$/Y; +/+; TH-Gal4/UAS-GtACR1 | Fed retinal, light surrounding threat, 41°C heat threat at 8 mins | 29 |
| B | CDN>GtACR1-EYFP | $w^{1118}$/Y; NP2719-Gal4, RepoGal80/+; UAS-GtACR1/+ | | |
| C | CDN>UAS-D2R | $w^{1118}$/Y; NP2719-Gal4, RepoGal80/+; UAS-Dop2R.S/+ | 41°C heat threat at 8 mins | 29 |
| | $D2R^{\Delta1}$, CDN-Gal4 | $D2R^{\Delta1}$/Y; NP2719-Gal4, RepoGal80/+; +/+ | 41°C heat threat at 8 mins | 27 |
| | $D2R^{\Delta1}$, UAS-D2R | $D2R^{\Delta1}$/Y; +/+; UAS-Dop2R.S/+ | 41°C heat threat at 8 mins | 26 |
| | $D2R^{\Delta1}$, CDN>UAS-D2R | $D2R^{\Delta1}$/Y; NP2719-Gal4, RepoGal80/+; UAS-Dop2R.S/+ | 41°C heat threat at 8 mins | 31 |
| D | CDN>GFP | $w^{1118}$, Dcr-2/Y; NP2719-Gal4, RepoGal80/+; UAS-myr::GFP/+ | 41°C heat threat at 8 mins | 29 |
| | D2R-RNAi/+ | $w^{1118}$/Y; +/+; UAS-D2R-RNAi-1/+ | 41°C heat threat at 8 mins | 25 |
| | CDN>D2R-RNAi-1 | $w^{1118}$, Dcr-2/Y; NP2719-Gal4, RepoGal80/+; UAS-D2R-RNAi-1/+ | 41°C heat threat at 8 mins | 36 |
| | D2R-RNAi-2/+ | $y^{1}sc^{*}v^{1}sev^{21}$/Y; +/+; UAS-D2R-RNAi-2/+ | 41°C heat threat at 8 mins | 26 |
| | CDN>D2R-RNAi-2 | $w^{1118}$, Dcr-2/Y; NP2719-Gal4, RepoGal80/+; UAS-UAS-D2R-RNAi-2/+ | 41°C heat threat at 8 mins | 44 |
| E | - | - | - | - |

| | | | | |
|---|---|---|---|---|
| F | CDN>ChR2-XXM, GCaMP6s | $w^{1118}$, Dcr-2/Y; NP2719-Gal4, RepoGal80/ChR2-XXM; UAS-GCaMP6s/+ | Saline | 10 |
| | CDN>ChR2-XXM, GCaMP6s | $w^{1118}$, Dcr-2/Y; NP2719-Gal4, RepoGal80/ChR2-XXM; UAS-GCaMP6s/+ | 100 $\mu$M DA | 10 |
| | CDN>ChR2-XXM, GCaMP6s + D2R-RNAi-1 | $w^{1118}$, Dcr-2/Y; NP2719-Gal4, RepoGal80/ChR2-XXM; UAS-GCaMP6s/UAS-D2R-RNAi-1 | Saline | 10 |
| | CDN>ChR2-XXM, GCaMP6s + D2R-RNAi-1 | $w^{1118}$, Dcr-2/Y; NP2719-Gal4, RepoGal80/ChR2-XXM; UAS-GCaMP6s/UAS-D2R-RNAi-1 | 100 $\mu$M DA | 10 |
| G | - | - | - | - |
| H | CDN>ChR2-XXM, green-camuiα | $w^{1118}$; NP2719-Gal4, RepoGal80/ChR2-XXM; UAS-green-camuiα/+ | Saline | 10 |
| | CDN>ChR2-XXM, green-camuiα | $w^{1118}$; NP2719-Gal4, RepoGal80/ChR2-XXM; UAS-green-camuiα/+ | 10 $\mu$M DA | 10 |
| | CDN>ChR2-XXM, green-camuiα +D2R-RNAi-1 | $w^{1118}$; NP2719-Gal4, RepoGal80/ChR2-XXM; UAS-green-camuiα/UAS-D2R-RNAi-1 | Saline | 10 |
| | CDN>ChR2-XXM, green-camuiα +D2R-RNAi-1 | $w^{1118}$; NP2719-Gal4, RepoGal80/ChR2-XXM; UAS-green-camuiα/UAS-D2R-RNAi-1 | 10 $\mu$M DA | 10 |
| **FIGURE 3** | | | | |
| A | TH>TrpA1, Not satiated | $w^{1118}$/Y; UAS-TrpA1/+; TH-Gal4/+ | Not satiated, tested at 28.5°C | 16 |
| | TH>TrpA1, Satiated | $w^{1118}$/Y; UAS-TrpA1/+; TH-Gal4/+ | Satiated 2.5 hrs at room temperature, tested at 28.5°C | 22 |
| B | CDN>ChR2-XXM, GCaMP6s | $w^{1118}$; NP2719-Gal4, RepoGal80/ChR2-XXM; UAS-GCaMP6s/+ | Saline, Not satiated | 20 |
| | CDN>ChR2-XXM, GCaMP6s | $w^{1118}$; NP2719-Gal4, RepoGal80/ChR2-XXM; UAS-GCaMP6s/+ | Saline, Satiated 4.5 hrs | 19 |
| | CDN>ChR2-XXM, GCaMP6s | $w^{1118}$; NP2719-Gal4, RepoGal80/ChR2-XXM; UAS-GCaMP6s/+ | 25 μM DA, Not satiated | 20 |
| | CDN>ChR2-XXM, GCaMP6s | $w^{1118}$; NP2719-Gal4, RepoGal80/ChR2-XXM; UAS-GCaMP6s/+ | 25 μM DA, Satiated 4.5 hrs | 17 |
| | CDN>ChR2-XXM, GCaMP6s | $w^{1118}$; NP2719-Gal4, RepoGal80/ChR2-XXM; UAS-GCaMP6s/+ | 50 μM DA, Not satiated | 20 |
| | CDN>ChR2-XXM, GCaMP6s | $w^{1118}$; NP2719-Gal4, RepoGal80/ChR2-XXM; UAS-GCaMP6s/+ | 50 μM DA, Satiated 4.5 hrs | 20 |

| | | | | |
|---|---|---|---|---|
| C | CDN>ChR2-XXM, green-camuiα | $w^{1118}$; NP2719-Gal4, RepoGal80/ChR2-XXM; UAS-GCaMP6s/+ | Saline, Not satiated | 8 |
| | CDN>ChR2-XXM, green-camuiα | $w^{1118}$; NP2719-Gal4, RepoGal80/ChR2-XXM; UAS-GCaMP6s/+ | Saline, Satiated 4.5 hrs | 10 |
| | CDN>ChR2-XXM, green-camuiα | $w^{1118}$; NP2719-Gal4, RepoGal80/ChR2-XXM; UAS-green-camuiα/+ | 10 µM DA, Not satiated | 19 |
| | CDN>ChR2-XXM, green-camuiα | $w^{1118}$; NP2719-Gal4, RepoGal80/ChR2-XXM; UAS-green-camuiα/+ | 10 µM DA, Satiated 4.5 hrs | 20 |
| **FIGURE 4** | | | | |
| A | - | - | - | - |
| B | CDN>GFP | $w^{1118}$, Dcr-2/Y; NP2719-Gal4, RepoGal80/+; UAS-mCD8::GFP/+ | 41°C heat threat at 8 mins | 33 |
| | UAS-$G_{\alpha o}$-RNAi-1/+ | $w^{1118}$/Y; +/+; UAS- UAS-$G_{\alpha o}$-RNAi-1 | 41°C heat threat at 8 mins | 30 |
| | CDN> UAS-$G_{\alpha o}$-RNAi-1 | $w^{1118}$, Dcr-2/Y; NP2719-Gal4, RepoGal80/+; UAS-mCD8::GFP/ UAS-$G_{\alpha o}$-RNAi-1 | 41°C heat threat at 8 mins | 25 |
| | UAS-$G_{\alpha o}$-RNAi-2/+ | $y^1sc^*v^1sev^{21}$/Y; +/+; UAS-$G_{\alpha o}$-RNAi-2/+ | 41°C heat threat at 8 mins | 33 |
| | CDN> UAS-$G_{\alpha o}$-RNAi-2 | $w^{1118}$, Dcr-2/Y; NP2719-Gal4, RepoGal80/+; UAS-mCD8::GFP/ UAS-$G_{\alpha o}$-RNAi-2 | 41°C heat threat at 8 mins | 30 |
| C | CDN>ChR2-XXM, GCaMP6s | $w^{1118}$, Dcr-2/Y; NP2719-Gal4, RepoGal80/ChR2-XXM; UAS-GCaMP6s/+ | Saline | 10 |
| | CDN>ChR2-XXM, GCaMP6s | $w^{1118}$, Dcr-2/Y; NP2719-Gal4, RepoGal80/ChR2-XXM; UAS-GCaMP6s/+ | 100 $\mu$M DA | 10 |
| | CDN>ChR2-XXM, GCaMP6s + UAS-$G_{\alpha o}$-RNAi-1 | $w^{1118}$, Dcr-2/Y; NP2719-Gal4, RepoGal80/ChR2-XXM; UAS-GCaMP6s/UAS- UAS-$G_{\alpha o}$-RNAi-1 | Saline | 10 |
| | CDN>ChR2-XXM, GCaMP6s + UAS-$G_{\alpha o}$-RNAi-1 | $w^{1118}$, Dcr-2/Y; NP2719-Gal4, RepoGal80/ChR2-XXM; UAS-GCaMP6s/UAS- UAS-$G_{\alpha o}$-RNAi-1 | 100 $\mu$M DA | 10 |
| D | - | - | - | - |
| E | CDN>eOPN3 | w+/Y; NP2719-Gal4, RepoGal80/+; UAS-eOPN3-HA/+ | Satiated, No light | 31 |
| | CDN>eOPN3 | w+/Y; NP2719-Gal4, RepoGal80/+; UAS-eOPN3-HA/+ | Satiated, Green light from mating start | 13 |
| F | CDN>ChR2-XXM, GCaMP6s | $w^{1118}$, Dcr-2/Y; NP2719-Gal4, RepoGal80/ChR2-XXM; UAS-GCaMP6s/+ | Not satiated | 10 |

| | | | | |
|---|---|---|---|---|
| | CDN>ChR2-XXM, GCaMP6s, eOPN3 | $w^{1118}$, Dcr-2/Y; NP2719-Gal4, RepoGal80/ChR2-XXM; UAS-GCaMP6s/UAS-eOPN3-HA | Not satiated | 10 |
| | CDN>ChR2-XXM, GCaMP6s | $w^{1118}$, Dcr-2/Y; NP2719-Gal4, RepoGal80/ChR2-XXM; UAS-GCaMP6s/+ | Satiated | 10 |
| | CDN>ChR2-XXM, GCaMP6s, eOPN3 | $w^{1118}$, Dcr-2/Y; NP2719-Gal4, RepoGal80/ChR2-XXM; UAS-GCaMP6s/UAS-eOPN3-HA | Satiated | 10 |
| G | CDN>eOPN3, No-prestimulation | w+/Y; NP2719-Gal4, RepoGal80/+; UAS-eOPN3-HA/+ | Spent 4.5 hrs in the dark prior to assay, 41°C heat threat at 8 mins | 29 |
| | CDN>eOPN3, Pre-stimulation | w+/Y; NP2719-Gal4, RepoGal80/+; UAS-eOPN3-HA/+ | Spent 4.5 hrs in green light prior to assay, 41°C heat threat at 8 mins | 30 |
| H | CDN>eOPN3, No-prestimulation | w+/Y; NP2719-Gal4, RepoGal80/+; UAS-eOPN3-HA/+ | Spent 4.5 hrs in the dark prior to assay, 41°C heat threat at 10 mins | 24 |
| | CDN>eOPN3, Pre-stimulation | w+/Y; NP2719-Gal4, RepoGal80/+; UAS-eOPN3-HA/+ | Spent 4.5 hrs in green light prior to assay, 41°C heat threat at 10 mins | 24 |
| **FIGURE 5** | | | | |
| A | - | - | - | - |
| B | CDN>GFP | $w^{1118}$, Dcr-2/Y; NP2719-Gal4, RepoGal80/+; UAS-myr::GFP/+ | Not satiated, 41°C heat threat at 8 mins | 25 |
| | CDN>GFP | $w^{1118}$, Dcr-2/Y; NP2719-Gal4, RepoGal80/+; UAS-myr::GFP/+ | Satiated 2.5 hrs, 41°C heat threat at 8 mins | 22 |
| | Krz-RNAi/+ | $w^{1118}$/Y; +/+; UAS-Krz-RNAi/+ | Not satiated, 41°C heat threat at 8 mins | 35 |
| | Krz-RNAi/+ | $w^{1118}$/Y; +/+; UAS-Krz-RNAi/+ | Satiated 2.5 hrs, 41°C heat threat at 8 mins | 27 |
| | CDN>Krz-RNAi | $w^{1118}$, Dcr-2/Y; NP2719-Gal4, RepoGal80/+; UAS-Krz-RNAi/+ | Not satiated, 41°C heat threat at 8 mins | 21 |
| | CDN>Krz-RNAi | $w^{1118}$, Dcr-2/Y; NP2719-Gal4, RepoGal80/+; UAS-Krz-RNAi/+ | Satiated 2.5 hrs, 41°C heat threat at 8 mins | 18 |
| C | CDN>ChR2-XXM, green-camuiα +Krz-RNAi | $w^{1118}$; NP2719-Gal4, RepoGal80/ChR2-XXM; UAS-green-camuiα/UAS-Krz-RNAi | Not satiated, 10 μM DA | 10 |
| | CDN>ChR2-XXM, green-camuiα +Krz-RNAi-1 | $w^{1118}$; NP2719-Gal4, RepoGal80/ChR2-XXM; | Satiated 4.5 hrs, 10 μM DA | 10 |

| | | | | |
|---|---|---|---|---|
| | | UAS-green-camuiα/UAS-Krz-RNAi | | |
| **FIGURE S1** | | | | |
| A | Canton-S | +/Y;+/+;+/+ | x hrs in satiety assay | 30 |
| B | CDN>GFP | $w^{1118}$, Dcr-2/Y; NP2719-Gal4, RepoGal80/+; UAS-mCD8::GFP/+ | 0 hrs in satiety assay, 41°C heat threat at 4 mins | 33 |
| | CDN>GFP | $w^{1118}$, Dcr-2/Y; NP2719-Gal4, RepoGal80/+; UAS-mCD8::GFP/+ | 1.5 hrs in satiety assay, 41°C heat threat at 4 mins | 20 |
| | CDN>GFP | $w^{1118}$, Dcr-2/Y; NP2719-Gal4, RepoGal80/+; UAS-mCD8::GFP/+ | 2.5 hrs in satiety assay, 41°C heat threat at 4 mins | 38 |
| | CDN>GFP | $w^{1118}$, Dcr-2/Y; NP2719-Gal4, RepoGal80/+; UAS-mCD8::GFP/+ | 3.5 hrs in satiety assay, 41°C heat threat at 4 mins | 20 |
| | CDN>GFP | $w^{1118}$, Dcr-2/Y; NP2719-Gal4, RepoGal80/+; UAS-mCD8::GFP/+ | 4.5 hrs in satiety assay, 41°C heat threat at 4 mins | 10 |
| C | Canton-S | +/Y;+/+;+/+ | 0 hrs in satiety assay | 25 |
| | Canton-S | +/Y;+/+;+/+ | 2.5 hrs in satiety assay | 16 |
| | Canton-S | +/Y;+/+;+/+ | 4.5 hrs in satiety assay | 21 |
| D | Males: CDN>GFP | $w^{1118}$, Dcr-2/Y; NP2719-Gal4, RepoGal80/+; UAS-myr::GFP/+ | Not satiated, 41°C heat threat at 8 mins | 23 |
| | Males: CDN>GFP | $w^{1118}$, Dcr-2/Y; NP2719-Gal4, RepoGal80/+; UAS-myr::GFP/+ | 2.5 hrs in satiety assay with $w^{1118}$ females, 41°C heat threat at 8 mins | 29 |
| | Males: CDN>GFP | $w^{1118}$, Dcr-2/Y; NP2719-Gal4, RepoGal80/+; UAS-myr-::GFP/+ | 2.5 hrs in satiety assay with Chr-mVenus females, 41°C heat threat at 8 mins | 25 |
| | Males: CDN>GFP | $w^{1118}$, Dcr-2/Y; NP2719-Gal4, RepoGal80/+; UAS-myr::GFP/+ | 2.5 hrs in satiety assay with Elav>SP females, 41°C heat threat at 8 mins | 30 |
| | Females: $w^{1118}$ | $w^{1118}$/ $w^{1118}$; +/+; +/+ | Used in satiety assay | |
| | Females: Chr-mVenus | $w^{1118}$, UAS-Chrimson-mVenus/$w^{1118}$, UAS-Chrimson-mVenus; +/+; +/+ | Used in satiety assay | |
| | Females: Elav>SP | $elav^{C155}/elav^{C155}$; UAS-Dicer2/+; UAS-SP/+ | Used in satiety assay | |
| E | - | - | - | - |
| F | Crz>TrpA1, No pre-stimulation | $w^{1118}$/Y; UAS-TrpA1/Crz-Gal4; +/+ | Spent 4.5 hrs at 23°C prior to assay | 25 |
| | Crz>TrpA1, Pre-stimulation | $w^{1118}$/Y; UAS-TrpA1/Crz-Gal4; +/+ | Spent 4.5 hrs at 28.5°C prior to assay | 24 |

| | | | | |
|---|---|---|---|---|
| G | Crz>TrpA1, No pre-stimulation | $w^{1118}$/Y; UAS-TrpA1/Crz-Gal4; +/+ | Spent 4.5 hrs at 23°C prior to assay, blue light stressor at 10 mins | 25 |
| | Crz>TrpA1, Pre-stimulation | $w^{1118}$/Y; UAS-TrpA1/Crz-Gal4; +/+ | Spent 4.5 hrs at 28.5°C prior to assay, blue light stressor at 10 mins | 24 |
| H | Crz>GtACR1, Not satiated | $w^{1118}$/Y; Crz-Gal4/+; UAS-GtACR1/+ | Not satiated | 20 |
| | Crz>GtACR1, Satiated (grey) | $w^{1118}$/Y; Crz-Gal4/+; UAS-GtACR1/+ | Satiated 2.5 hrs in dim light | 20 |
| | Crz>GtACR1, Satiated (green +Light) | $w^{1118}$/Y; Crz-Gal4/+; UAS-GtACR1/+ | Satiated 2.5 hrs in green light | 20 |
| I | Crz>GtACR1, Not satiated | $w^{1118}$/Y; Crz-Gal4/+; UAS-GtACR1/+ | Not satiated, 41°C heat threat at 8 mins | 27 |
| | Crz>GtACR1, Satiated (grey) | $w^{1118}$/Y; Crz-Gal4/+; UAS-GtACR1/+ | Satiated 2.5 hrs in dim light, 41°C heat threat at 8 mins | 30 |
| | Crz>GtACR1, Satiated (green +Light) | $w^{1118}$/Y; Crz-Gal4/+; UAS-GtACR1/+ | Satiated 2.5 hrs in green light, 41°C heat threat at 8 mins | 29 |
| **FIGURE S2** | | | | |
| A | - | - | - | - |
| B | - | - | - | - |
| C | UAS-TrpA1/+, 23°C pre-treatment | $w^{1118}$/Y; UAS-TrpA1/+; +/+ | Spent 4.5 hrs at 23°C prior to assay | |
| | UAS-TrpA1/+, 30°C pre-treatment | $w^{1118}$/Y; UAS-TrpA1/+; +/+ | Spent 4.5 hrs at 30°C prior to assay | |
| | P1>TrpA1, Not pre-stimulated | $w^{1118}$/Y; UAS-TrpA1/+; P1-Gal4/+ | Spent 4.5 hrs at 23°C prior to assay, blue light stressor at 10 min | 31 |
| | P1>TrpA1, Pre-stimulated | $w^{1118}$/Y; UAS-TrpA1/+; P1-Gal4/+ | Spent 4.5 hrs at 30°C prior to assay, blue light stressor at 10 min | 29 |
| | NPF>TrpA1, Not pre-stimulated | $w^{1118}$/Y; UAS-TrpA1/+; NPF-Gal4/+ | Spent 4.5 hrs at 23°C prior to assay, blue light stressor at 10 min | 25 |
| | NPF>TrpA1, Pre-stimulated | $w^{1118}$/Y; UAS-TrpA1/+; NPF-Gal4/+ | Spent 4.5 hrs at 30°C prior to assay, blue light stressor at 10 min | 31 |
| | pCd>TrpA1, Not pre-stimulated | $w^{1118}$/Y; UAS-TrpA1/+; pCd-Gal4/+ | Spent 4.5 hrs at 23°C prior to assay, blue light stressor at 10 min | 28 |
| | pCd>TrpA1, Pre-stimulated | $w^{1118}$/Y; UAS-TrpA1/+; pCd-Gal4/+ | Spent 4.5 hrs at 30°C prior to assay, blue light stressor at 10 min | 36 |

| | | | | |
|---|---|---|---|---|
| | NPF,pCd>TrpA1, Not pre-stimulated | $w^{1118}$/Y; UAS-TrpA1/+; pCd-Gal4/NPF-Gal4 | Spent 4.5 hrs at 23°C prior to assay, blue light stressor at 10 min | 27 |
| | NPF,pCd>TrpA1, Pre-stimulated | $w^{1118}$/Y; UAS-TrpA1/+; pCd-Gal4/NPF-Gal4 | Spent 4.5 hrs at 30°C prior to assay, blue light stressor at 10 min | 30 |
| | mAL>TrpA1, Not pre-stimulated | $w^{1118}$/Y; UAS-TrpA1/+; mAL-Gal4/+ | Spent 4.5 hrs at 23°C prior to assay, blue light stressor at 10 min | 29 |
| | mAL>TrpA1, Pre-stimulated | $w^{1118}$/Y; UAS-TrpA1/+; mAL-Gal4/+ | Spent 4.5 hrs at 30°C prior to assay, blue light stressor at 10 min | 34 |
| | vAB3>TrpA1, Not pre-stimulated | $w^{1118}$/Y; UAS-TrpA1/+; vAB3-Gal4/+ | Spent 4.5 hrs at 23°C prior to assay, blue light stressor at 10 min | 27 |
| | vAB3>TrpA1, Pre-stimulated | $w^{1118}$/Y; UAS-TrpA1/+; vAB3-Gal4/+ | Spent 4.5 hrs at 30°C prior to assay, blue light stressor at 10 min | 7 |
| | PPN1>TrpA1, Not pre-stimulated | $w^{1118}$/Y; UAS-TrpA1/+; PPN1-Gal4/+ | Spent 4.5 hrs at 23°C prior to assay, blue light stressor at 10 min | 26 |
| | PPN1>TrpA1, Pre-stimulated | $w^{1118}$/Y; UAS-TrpA1/+; PPN1-Gal4/+ | Spent 4.5 hrs at 30°C prior to assay, blue light stressor at 10 min | 31 |
| | NP5945>TrpA1, Not pre-stimulated | $w^{1118}$/Y; UAS-TrpA1/+; NP5945/+ | Spent 4.5 hrs at 23°C prior to assay, blue light stressor at 10 min | 30 |
| | NP5945>TrpA1, Pre-stimulated | $w^{1118}$/Y; UAS-TrpA1/+; NP5945/+ | Spent 4.5 hrs at 28.5°C prior to assay, blue light stressor at 10 min | 32 |
| | VT02857>TrpA1, Not pre-stimulated | $w^{1118}$/Y; UAS-TrpA1/+; VT02857/+ | Spent 4.5 hrs at 23°C prior to assay, blue light stressor at 10 min | 40 |
| | VT02857>TrpA1, Pre-stimulated | $w^{1118}$/Y; UAS-TrpA1/+; VT02857/+ | Spent 4.5 hrs at 28.5°C prior to assay, blue light stressor at 10 min | 36 |
| | VT02857,NP5945>TrpA1, Not pre-stimulated | $w^{1118}$/Y; UAS-TrpA1/+; NP5945/VT02857 | Spent 4.5 hrs at 23°C prior to assay, blue light stressor at 10 min | 33 |
| | VT02857,NP5945>TrpA1, Pre-stimulated | $w^{1118}$/Y; UAS-TrpA1/+; NP5945/VT02857 | Spent 4.5 hrs at 28.5°C prior to assay, blue light stressor at 10 min | 37 |

| | | | | |
|---|---|---|---|---|
| D | CRN>TrpA1, Not pre-stimulated | $w^{1118}$/Y; UAS-TrpA1/CRN-Gal4; +/+ | Spent 4.5 hrs at 23°C prior to assay, blue light stressor at 10 min | 31 |
| | CRN>TrpA1, pre-stimulated | $w^{1118}$/Y; UAS-TrpA1/CRN-Gal4; +/+ | Spent 4.5 hrs at 30°C prior to assay, blue light stressor at 10 min | 33 |
| E | CRN>TrpA1, Not pre-stimulated | $w^{1118}$/Y; UAS-TrpA1/CRN-Gal4; +/+ | Spent 4.5 hrs at 23°C prior to assay | 27 |
| | CRN>TrpA1, pre-stimulated | $w^{1118}$/Y; UAS-TrpA1/CRN-Gal4; +/+ | Spent 4.5 hrs at 30°C prior to assay | 27 |
| **FIGURE S3** | | | | |
| A | TH>GFP, Not stimulated | $w^{1118}$/Y; +/+; TH-Gal4/UAS-myrGFP | Spent 4.5 hrs at 23°C prior to assay | 25 |
| | TH>GFP, Pre-stimulated | $w^{1118}$/Y; +/+; TH-Gal4/UAS-myrGFP | Spent 4.5 hrs at 28.5°C prior to assay | 26 |
| | TrpA1/+, Not stimulated | $w^{1118}$/Y; UAS-TrpA1/+; +/+ | Spent 4.5 hrs at 23°C prior to assay | 32 |
| | TrpA1/+, Pre-stimulated | $w^{1118}$/Y; UAS-TrpA1/+; +/+ | Spent 4.5 hrs at 28.5°C prior to assay | 33 |
| B | TH>GtACR1-EYFP | $w^{1118}$/Y; +/+; TH-Gal4/UAS-GtACR1 | | |
| | TH>GtACR1-EYFP + Tsh-Gal80 | $w^{1118}$/Y; Tsh-Gal80/+; TH-Gal4/UAS-GtACR1 | | |
| C | TH>GFP, Not stimulated | $w^{1118}$/Y; +/+; TH-Gal4/UAS-myrGFP | Spent 4.5 hrs at 23°C prior to assay | 17 |
| | TH>GFP, Pre-stimulated | $w^{1118}$/Y; +/+; TH-Gal4/UAS-myrGFP | Spent 4.5 hrs at 28.5°C prior to assay | 14 |
| | TrpA1/+, Not stimulated | $w^{1118}$/Y; UAS-TrpA1/+; +/+ | Spent 4.5 hrs at 23°C prior to assay | 16 |
| | TrpA1/+, Pre-stimulated | $w^{1118}$/Y; UAS-TrpA1/+; +/+ | Spent 4.5 hrs at 28.5°C prior to assay | 17 |
| | TH>TrpA1, Not stimulated | $w^{1118}$/Y; UAS-TrpA1/+; TH-Gal4/+ | Spent 4.5 hrs at 23°C prior to assay | 14 |
| | TH>TrpA1, Pre-stimulated | $w^{1118}$/Y; UAS-TrpA1/+; TH-Gal4/+ | Spent 4.5 hrs at 28.5°C prior to assay | 15 |
| | TH>TrpA1 + Tsh-Gal80, Not stimulated | $w^{1118}$/Y; UAS-TrpA1/Tsh-Gal80; TH-Gal4/+ | Spent 4.5 hrs at 23°C prior to assay | 17 |
| | TH>TrpA1 + Tsh-Gal80, Pre-stimulated | $w^{1118}$/Y; UAS-TrpA1/Tsh-Gal80; TH-Gal4/+ | Spent 4.5 hrs at 28.5°C prior to assay | 15 |
| D | TH>GFP, 23°C | $w^{1118}$/Y; +/+; TH-Gal4/UAS-myrGFP | Assay at 23°C | 19 |
| | TH>GFP, 28.5°C | $w^{1118}$/Y; +/+; TH-Gal4/UAS-myrGFP | Assay at 28.5°C | 17 |
| | TrpA1/+, 23°C | $w^{1118}$/Y; UAS-TrpA1/+; +/+ | Assay at 23°C | 22 |
| | TrpA1/+, 28.5°C | $w^{1118}$/Y; UAS-TrpA1/+; +/+ | Assay at 28.5°C | 17 |

| | | | | |
|---|---|---|---|---|
| | TH>TrpA1, 23°C | $w^{1118}$/Y; UAS-TrpA1/+; TH-Gal4/+ | Assay at 23°C | 21 |
| | TH>TrpA1, 28.5°C | $w^{1118}$/Y; UAS-TrpA1/+; TH-Gal4/+ | Assay at 28.5°C | 21 |
| E | TH>TrpA1, 23°C | $w^{1118}$/Y; UAS-TrpA1/+; TH-Gal4/+ | Assay at 23°C, 41°C heat threat at 8 mins | 38 |
| | TH>TrpA1, 28.5°C | $w^{1118}$/Y; UAS-TrpA1/+; TH-Gal4/+ | Assay at 28.5°C, 41°C heat threat at 8 mins | 15 |
| F | TH>GtACR1-EYFP, No light | $w^{1118}$/Y; +/+; TH-Gal4/UAS-GtACR1 | Spent 2.5 hrs in a satiety assay in dim light | 13 |
| | TH>GtACR1-EYFP, Light | $w^{1118}$/Y; +/+; TH-Gal4/UAS-GtACR1 | Spent 2.5 hrs in a satiety assay in green light | 25 |
| G | TH>GtACR1, no light | $w^{1118}$/Y; +/+; TH-Gal4/UAS-GtACR1 | Fed retinal, but no light during testing, | 23 |
| | TH>GtACR1, no retinal | $w^{1118}$/Y; +/+; TH-Gal4/UAS-GtACR1 | Not fed retinal, green light during testing | 20 |
| | TH>GtACR1 | $w^{1118}$/Y; +/+; TH-Gal4/UAS-GtACR1 | Fed retinal, green light during testing | 18 |
| H | TH>GtACR1-EYFP, No light | $w^{1118}$/Y; +/+; TH-Gal4/UAS-GtACR1 | Spent 2.5 hrs in a satiety assay in dim light, then spent 200 mins in dim light, recovered 24 hrs | 8 |
| | TH>GtACR1-EYFP, Light | $w^{1118}$/Y; +/+; TH-Gal4/UAS-GtACR1 | Spent 2.5 hrs in a satiety assay in dim light, then spent 20 min on/20 min off of green light for 200 mins (pattern repeated 5x), recovered 24 hrs | 10 |
| **FIGURE S4** | | | | |
| A | WT | Canton-S | 41°C heat threat at 8 mins | 28 |
| | DopEcR | $w^{1118}$/Y; DopEcR/DopEcR; +/+ | 41°C heat threat at 8 mins | 28 |
| | DopR1 | +/Y;+/+; $DopR1^{attp}$/ $DopR1^{attp}$ | 41°C heat threat at 8 mins | 33 |
| | DopR2 | +/Y;+/+; $DopR2^{attp}$/$DopR2^{attp}$ | 41°C heat threat at 8 mins | 28 |
| | D2R | $D2R^{\Delta 1}$/Y; +/+; +/+ | 41°C heat threat at 8 mins | 27 |
| C | WT | Canton-S | | 10 |
| | DopEcR | $w^{1118}$/Y; DopEcR/DopEcR; +/+ | | 15 |
| | DopR1 | +/Y;+/+; $DopR1^{attp}$/ $DopR1^{attp}$ | | 10 |
| | DopR2 | +/Y;+/+; $DopR2^{attp}$/$DopR2^{attp}$ | | 12 |
| | D2R | $D2R^{\Delta 1}$/Y; +/+; +/+ | | 11 |

| | | | | |
|---|---|---|---|---|
| D | TH>GtACR1-EYFP, No light | $w^{1118}$/Y; +/+; TH-Gal4/UAS-GtACR1 | No light, 41°C heat threat at 8 mins | 27 |
| | TH>GtACR1-EYFP, Light | $w^{1118}$/Y; +/+; TH-Gal4/UAS-GtACR1 | Green light 7.5 to 9.5 mins into mating, 41°C heat threat at 8 mins | 33 |
| | TH, CDN>GtACR1-EYFP, No light | $w^{1118}$/Y; NP2719-Gal4, RepoGal80/+; TH-Gal4/UAS-GtACR1 | No light, 41°C heat threat at 8 mins | 26 |
| | TH, CDN>GtACR1-EYFP, Light | $w^{1118}$/Y; NP2719-Gal4, RepoGal80/+; TH-Gal4/UAS-GtACR1 | Green light 7.5 to 9.5 mins into mating, 41°C heat threat at 8 mins | 26 |
| E | CDN>UAS-D2R | $w^{1118}$/Y; NP2719-Gal4, RepoGal80/+; UAS-Dop2R.S/+ | | 19 |
| | $D2R^{\Delta 1}$, CDN-Gal4 | $D2R^{\Delta 1}$/Y; NP2719-Gal4, RepoGal80/+; +/+ | | 22 |
| | $D2R^{\Delta 1}$, UAS-D2R | $D2R^{\Delta 1}$/Y; +/+; UAS-Dop2R.S/+ | | 36 |
| | $D2R^{\Delta 1}$, CDN>UAS-D2R | $D2R^{\Delta 1}$/Y; NP2719-Gal4, RepoGal80/+; UAS-Dop2R.S/+ | | 23 |
| F | CDN>GFP | $w^{1118}$/Y; NP2719-Gal4, RepoGal80/+; UAS-myr::GFP/+ | 41°C heat threat at 10 mins | 32 |
| | UAS-D2R/+ | $w^{1118}$/Y; +/+; UAS-Dop2R.S/+ | 41°C heat threat at 10 mins | 27 |
| | CDN>UAS-D2R | $w^{1118}$/Y; NP2719-Gal4, RepoGal80/+; UAS-Dop2R.S/+ | 41°C heat threat at 10 mins | 28 |
| G | CDN>GFP | $w^{1118}$, Dcr-2/Y; NP2719-Gal4, RepoGal80/+; UAS-myr::GFP/+ | | 16 |
| | D2R-RNAi/+ | $w^{1118}$/Y; +/+; UAS-D2R-RNAi-1/+ | | 12 |
| | CDN>D2R-RNAi-1 | $w^{1118}$, Dcr-2/Y; NP2719-Gal4, RepoGal80/+; UAS-D2R-RNAi-1/+ | | 13 |
| | D2R-RNAi-2/+ | $y^{1}sc^{*}v^{1}sev^{21}$/Y; +/+; UAS-D2R-RNAi-2/+ | | 12 |
| | CDN>D2R-RNAi-2 | $w^{1118}$, Dcr-2/Y; NP2719-Gal4, RepoGal80/+; UAS-UAS-D2R-RNAi-2/+ | | 17 |
| **FIGURE S5** | | | | |
| A | CDN>GCaMP6s | $w^{1118}$, Dcr-2/Y; NP2719-Gal4, RepoGal80; UAS-GCaMP6s/+ | Saline | 10 |
| | CDN>GCaMP6s | $w^{1118}$, Dcr-2/Y; NP2719-Gal4, RepoGal80; UAS-GCaMP6s/+ | 100 μM DA | 10 |
| B | CDN>ChR2-XXM, GCaMP6s | $w^{1118}$, Dcr-2/Y; NP2719-Gal4, RepoGal80/ChR2-XXM; UAS-GCaMP6s/+ | No perfusion | 10 |

| | | | | |
|---|---|---|---|---|
| C | CDN>ChR2-XXM, GCaMP6s | $w^{1118}$, Dcr-2/Y; NP2719-Gal4, RepoGal80/ChR2-XXM; UAS-GCaMP6s/+ | Saline | 10 |
| | CDN>ChR2-XXM, GCaMP6s | $w^{1118}$, Dcr-2/Y; NP2719-Gal4, RepoGal80/ChR2-XXM; UAS-GCaMP6s/+ | 100 $\mu$M DA | 10 |
| D | D2R$^{\Delta}$1, CDN>ChR2-XXM, GCaMP6s | D2R$^{\Delta}$1/Y; NP2719-Gal4, RepoGal80/ChR2-XXM; UAS-GCaMP6s/+ | Saline | 10 |
| | D2R$^{\Delta}$1, CDN>ChR2-XXM, GCaMP6s | D2R$^{\Delta}$1/Y; NP2719-Gal4, RepoGal80/ChR2-XXM; UAS-GCaMP6s/+ | 100 $\mu$M DA | 10 |
| E | CDN>ChR2-XXM, GCaMP6s + D2R-RNAi-1 | $w^{1118}$, Dcr-2/Y; NP2719-Gal4, RepoGal80/ChR2-XXM; UAS-GCaMP6s/UAS-D2R-RNAi-1 | Saline | 10 |
| | CDN>ChR2-XXM, GCaMP6s + D2R-RNAi-1 | $w^{1118}$, Dcr-2/Y; NP2719-Gal4, RepoGal80/ChR2-XXM; UAS-GCaMP6s/UAS-D2R-RNAi-1 | 100 $\mu$M DA | 10 |
| F | CDN>ChR2-XXM, GCaMP6s + D2R-RNAi-2 | $w^{1118}$, Dcr-2/Y; NP2719-Gal4, RepoGal80/ChR2-XXM; UAS-GCaMP6s/UAS-D2R-RNAi-2 | Saline | 10 |
| | CDN>ChR2-XXM, GCaMP6s + D2R-RNAi-2 | $w^{1118}$, Dcr-2/Y; NP2719-Gal4, RepoGal80/ChR2-XXM; UAS-GCaMP6s/UAS-D2R-RNAi-2 | 100 $\mu$M DA | 10 |
| G | CDN>green-camuiα | $w^{1118}$; NP2719-Gal4, RepoGal80/ChR2-XXM; UAS-green-camuiα/UAS-D2R-RNAi-2 | 10 μM DA | 10 |
| H | D2R$^{\Delta 1}$, CDN>ChR2-XXM, green-camuiα | $w^{1118}$; NP2719-Gal4, RepoGal80/ChR2-XXM; UAS-green-camuiα/UAS-D2R-RNAi-2 | Saline | 10 |
| | D2R$^{\Delta 1}$, CDN>ChR2-XXM, green-camuiα | $w^{1118}$; NP2719-Gal4, RepoGal80/ChR2-XXM; UAS-green-camuiα/UAS-D2R-RNAi-2 | 10 $\mu$M DA | 10 |
| I | CDN>ChR2-XXM, green-camuiα +D2R-RNAi-2 | $w^{1118}$; NP2719-Gal4, RepoGal80/ChR2-XXM; UAS-green-camuiα/UAS-D2R-RNAi-2 | Saline | 10 |
| | CDN>ChR2-XXM, green-camuiα +D2R-RNAi-2 | $w^{1118}$; NP2719-Gal4, RepoGal80/ChR2-XXM; UAS-green-camuiα/UAS-D2R-RNAi-2 | 10 $\mu$M DA | 10 |

| FIGURE S6 | | | | |
|---|---|---|---|---|
| A | CDN>GtACR1, Not satiated, No light | $w^{1118}$/Y; NP2719-Gal4, RepoGal80/+; UAS-GtACR1/+ | Not satiated, tested in the dark, 41°C heat threat at 8 min | 23 |
| | CDN>GtACR1, Not satiated, Light | $w^{1118}$/Y; NP2719-Gal4, RepoGal80/+; UAS-GtACR1/+ | Not satiated, green light from mating start, 41°C heat threat at 8 min | 22 |
| | CDN>GtACR1, Satiated, No light | $w^{1118}$/Y; NP2719-Gal4, RepoGal80/+; UAS-GtACR1/+ | Satiated 2.5 hrs in dim light, tested in the dark, 41°C heat threat at 8 min | 16 |
| | CDN>GtACR1, Satiated, Light | $w^{1118}$/Y; NP2719-Gal4, RepoGal80/+; UAS-GtACR1/+ | Satiated 2.5 hrs in dim light, green light from mating start, 41°C heat threat at 8 min | 20 |
| B | CDN>tdTomato, GCaMP6s, Not satiated | $w^{1118}$/Y; NP2719-Gal4, RepoGal80/UAS-opGCaMP6s; UAS-myr::tdTomato/+ | Not satiated | 10 |
| | CDN>tdTomato, GCaMP6s, Satiated | $w^{1118}$/Y; NP2719-Gal4, RepoGal80/UAS-opGCaMP6s; UAS-myr::tdTomato/+ | Satiated 4.5 hrs | 10 |
| C | CDN>Chr, Not satiated, Green light | +/Y; NP2719-Gal4, RepoGal80/+; UAS-Chrimson/+ | Fed retinal, Not satiated, 1 min of 10 μW/mm$^2$ green light at 8 min | 24 |
| | CDN>Chr, Satiated, Green light | +/Y; NP2719-Gal4, RepoGal80/+; UAS-Chrimson/+ | Fed retinal, Satiated 2.5 hrs in dim light, 1 min of 10 μW/mm$^2$ green light at 8 min | 26 |
| | CDN>Chr, Not satiated, 39°C heat | +/Y; NP2719-Gal4, RepoGal80/+; UAS-Chrimson/+ | Fed retinal, Not satiated, 39°C heat threat at 8 min | 23 |
| | CDN>Chr, Satiated, 39°C heat | +/Y; NP2719-Gal4, RepoGal80/+; UAS-Chrimson/+ | Fed retinal, Satiated 2.5 hrs, 39°C heat threat at 8 min | 22 |
| D | CDN>Chr, Not satiated, Green light | +/Y; NP2719-Gal4, RepoGal80/+; UAS-Chrimson/+ | Fed retinal, Not satiated, 1 min of 15 μW/mm$^2$ green light at 8 min | 32 |
| | CDN>Chr, Satiated, Green light | +/Y; NP2719-Gal4, RepoGal80/+; UAS-Chrimson/+ | Fed retinal, Satiated 2.5 hrs in dim light, 1 min of 15 μW/mm$^2$ green light at 8 min | 34 |
| | CDN>Chr, Not satiated, 41°C heat | +/Y; NP2719-Gal4, RepoGal80/+; UAS-Chrimson/+ | Fed retinal, Not satiated, 41°C heat threat at 8 min | 25 |
| | CDN>Chr, Satiated, 41°C heat | +/Y; NP2719-Gal4, RepoGal80/+; UAS-Chrimson/+ | Fed retinal, Satiated 2.5 hrs, | 26 |

| | | | | |
|---|---|---|---|---|
| | | | 41°C heat threat at 8 min | |
| E | CDN>Chr, Not satiated | $w^{1118}$, Dcr-2/Y; NP2719-Gal4, RepoGal80/+; UAS-Chrimson/+ | Fed retinal, Not satiated, 1 min of 15 µW/mm$^2$ green light at 8 min | 26 |
| | CDN>Chr, Satiated | $w^{1118}$, Dcr-2/Y; NP2719-Gal4, RepoGal80/+; UAS-Chrimson/+ | Fed retinal, Satiated 2.5 hrs in dim light, 1 min of 15 µW/mm$^2$ green light at 8 min | 25 |
| | CDN>Chr + D2R-RNAi-2 | $w^{1118}$, Dcr-2/Y; NP2719-Gal4, RepoGal80/+; UAS-Chrimson/UAS-D2R-RNAi-2 | Fed retinal, Not satiated, 1 min of 15 µW/mm$^2$ green light at 8 min | 33 |
| **FIGURE S7** | | | | |
| A | TH>TrpA1, 0 hrs satiated | $w^{1118}$/Y; UAS-TrpA1/+; TH-Gal4/+ | Not satiated, Assay at 28.5°C | 18 |
| | TH>TrpA1, 1.5 hrs satiated | $w^{1118}$/Y; UAS-TrpA1/+; TH-Gal4/+ | Satiated 1.5 hrs at 23°C, Assay at 28.5°C | 21 |
| | TH>TrpA1, 2.5 hrs satiated (reproduced from **Figure 3A**) | $w^{1118}$/Y; UAS-TrpA1/+; TH-Gal4/+ | Satiated 2.5 hrs at 23°C, Assay at 28.5°C | 21 |
| | TH>TrpA1, 3.5 hrs satiated | $w^{1118}$/Y; UAS-TrpA1/+; TH-Gal4/+ | Satiated 3.5 hrs at 23°C, Assay at 28.5°C | 19 |
| | TH>TrpA1, 4.5 hrs satiated | $w^{1118}$/Y; UAS-TrpA1/+; TH-Gal4/+ | Satiated 4.5 hrs at 23°C, Assay at 28.5°C | 18 |
| B | TH>tdTomato, GCaMP6s, Not satiated | $w^{1118}$/Y; UAS-opGCaMP6s/+; TH-Gal4/UAS-myr::tdTomato | Not satiated | 6 |
| | TH>tdTomato, GCaMP6s, Satiated | $w^{1118}$/Y; UAS-opGCaMP6s/+; TH-Gal4/UAS-myr::tdTomato | Satiated 4.5 hrs | 9 |
| C | TH>ChR2-XXM, CGaMP6s, Not Satiated | $w^{1118}$/Y; UAS-ChR2-XXM/+; TH-Gal4/UAS-GCaMP6s | Not satiated, 1 second blue light pulse | 4 |
| | TH>ChR2-XXM, CGaMP6s, Satiated | $w^{1118}$/Y; UAS-ChR2-XXM/+; TH-Gal4/UAS-GCaMP6s | Satiated 4.5 hrs, 1 second blue light pulse | 6 |
| D | CDN>ChR2-XXM, GCaMP6s | $w^{1118}$; NP2719-Gal4, RepoGal80/ChR2-XXM; UAS-GCaMP6s/+ | Saline, Not satiated | 20 |
| | CDN>ChR2-XXM, GCaMP6s | $w^{1118}$; NP2719-Gal4, RepoGal80/ChR2-XXM; UAS-GCaMP6s/+ | Saline, Satiated 4.5 hrs | 19 |
| E | CDN>ChR2-XXM, GCaMP6s | $w^{1118}$; NP2719-Gal4, RepoGal80/ChR2-XXM; UAS-GCaMP6s/+ | 25 µM DA, Not satiated | 20 |
| | CDN>ChR2-XXM, GCaMP6s | $w^{1118}$; NP2719-Gal4, RepoGal80/ChR2-XXM; UAS-GCaMP6s/+ | 25 µM DA, Satiated 4.5 hrs | 17 |

| | | | | |
|---|---|---|---|---|
| F | CDN>ChR2-XXM, GCaMP6s | $w^{1118}$; NP2719-Gal4, RepoGal80/ChR2-XXM; UAS-GCaMP6s/+ | 50 µM DA, Not satiated | 20 |
| | CDN>ChR2-XXM, GCaMP6s | $w^{1118}$; NP2719-Gal4, RepoGal80/ChR2-XXM; UAS-GCaMP6s/+ | 50 µM DA, Satiated 4.5 hrs | 20 |
| G | CDN>ChR2-XXM, GCaMP6s | $w^{1118}$; NP2719-Gal4, RepoGal80/ChR2-XXM; UAS-GCaMP6s/+ | 100 µM DA, Not satiated | 10 |
| | CDN>ChR2-XXM, GCaMP6s | $w^{1118}$; NP2719-Gal4, RepoGal80/ChR2-XXM; UAS-GCaMP6s/+ | 100 µM DA, Satiated 4.5 hrs | 10 |
| H | CDN>ChR2-XXM, green-camuiα | $w^{1118}$; NP2719-Gal4, RepoGal80/ChR2-XXM; UAS-GCaMP6s/+ | Saline, Not satiated | 8 |
| | CDN>ChR2-XXM, green-camuiα | $w^{1118}$; NP2719-Gal4, RepoGal80/ChR2-XXM; UAS-GCaMP6s/+ | Saline, Satiated 4.5 hrs | 10 |
| I | CDN>ChR2-XXM, green-camuiα | $w^{1118}$; NP2719-Gal4, RepoGal80/ChR2-XXM; UAS-green-camuiα/+ | 10 *µ*M DA, Not satiated | 10 |
| | CDN>ChR2-XXM, green-camuiα | $w^{1118}$; NP2719-Gal4, RepoGal80/ChR2-XXM; UAS-green-camuiα/+ | 10 *µ*M DA, Not satiated | 10 |
| **FIGURE S8** | | | | |
| A | CDN>GFP | $w^{1118}$, Dcr-2/Y; NP2719-Gal4, RepoGal80/+; UAS-myr::GFP/+ | 41°C heat threat at 8 mins | 18 |
| | UAS-$G_{\alpha o}$-RNAi-1/+ | $w^{1118}$/Y; +/+; UAS- UAS-$G_{\alpha o}$-RNAi-1 | 41°C heat threat at 8 mins | 16 |
| | CDN> UAS-$G_{\alpha o}$-RNAi-1 | $w^{1118}$, Dcr-2/Y; NP2719-Gal4, RepoGal80/+; UAS-$G_{\alpha o}$-RNAi-1/+ | 41°C heat threat at 8 mins | 14 |
| | UAS-$G_{\alpha o}$-RNAi-2/+ | $y^{1}sc^{*}v^{1}sev^{21}$/Y; +/+; UAS-$G_{\alpha o}$-RNAi-2/+ | 41°C heat threat at 8 mins | 15 |
| | CDN> UAS-$G_{\alpha o}$-RNAi-2 | $w^{1118}$, Dcr-2/Y; NP2719-Gal4, RepoGal80/+; UAS-$G_{\alpha o}$-RNAi-2/+ | 41°C heat threat at 8 mins | 15 |
| B | CDN>ChR2-XXM, GCaMP6s | $w^{1118}$, Dcr-2/Y; NP2719-Gal4, RepoGal80/ChR2-XXM; UAS-GCaMP6s/+ | Saline | 10 |
| | CDN>ChR2-XXM, GCaMP6s | $w^{1118}$, Dcr-2/Y; NP2719-Gal4, RepoGal80/ChR2-XXM; UAS-GCaMP6s/+ | 100 *µ*M DA | 10 |
| C | CDN>ChR2-XXM, GCaMP6s + UAS-$G_{\alpha o}$-RNAi-1 | $w^{1118}$, Dcr-2/Y; NP2719-Gal4, RepoGal80/ChR2-XXM; UAS-GCaMP6s/UAS- UAS-$G_{\alpha o}$-RNAi-1 | Saline | 10 |
| | CDN>ChR2-XXM, GCaMP6s + UAS-$G_{\alpha o}$-RNAi-1 | $w^{1118}$, Dcr-2/Y; NP2719-Gal4, RepoGal80/ChR2-XXM; UAS-GCaMP6s/UAS- UAS-$G_{\alpha o}$-RNAi-1 | 100 *µ*M DA | 10 |

| | | | | |
|---|---|---|---|---|
| D | CDN>eOPN3 | w+/Y; NP2719-Gal4, RepoGal80/+; UAS-eOPN3-HA/+ | No light | 26 |
| | CDN>eOPN3 | w+/Y; NP2719-Gal4, RepoGal80/+; UAS-eOPN3-HA/+ | Satiated, Green light from mating start | 24 |
| **FIGURE S9** | | | | |
| A | Krz-RNAi/+, 4 min | $w^{1118}$/Y; +/+; UAS-Krz-RNAi/+ | 41°C heat threat at 4 mins | 25 |
| | Krz-RNAi/+, 8 min | $w^{1118}$/Y; +/+; UAS-Krz-RNAi/+ | 41°C heat threat at 8 mins | 33 |
| | Krz-RNAi/+, 12 min | $w^{1118}$/Y; +/+; UAS-Krz-RNAi/+ | 41°C heat threat at 12 mins | 25 |
| | Krz-RNAi/+, 16 min | $w^{1118}$/Y; +/+; UAS-Krz-RNAi/+ | 41°C heat threat at 16 mins | 24 |
| | CDN>GFP, 4 min | $w^{1118}$, Dcr-2/Y; NP2719-Gal4, RepoGal80/+; UAS-CD8::GFP/+ | 41°C heat threat at 4 mins | 25 |
| | CDN>GFP, 8 min | $w^{1118}$, Dcr-2/Y; NP2719-Gal4, RepoGal80/+; UAS-CD8::GFP/+ | 41°C heat threat at 8 mins | 29 |
| | CDN>GFP, 12 min | $w^{1118}$, Dcr-2/Y; NP2719-Gal4, RepoGal80/+; UAS-CD8::GFP/+ | 41°C heat threat at 12 mins | 29 |
| | CDN>GFP, 16 min | $w^{1118}$, Dcr-2/Y; NP2719-Gal4, RepoGal80/+; UAS-CD8::GFP/+ | 41°C heat threat at 16 mins | 30 |
| | CDN>Krz-RNAi, 4 min | $w^{1118}$, Dcr-2/Y; NP2719-Gal4, RepoGal80/+; UAS-Krz-RNAi/+ | 41°C heat threat at 4 mins | 20 |
| | CDN>Krz-RNAi, 8 min | $w^{1118}$, Dcr-2/Y; NP2719-Gal4, RepoGal80/+; UAS-Krz-RNAi/+ | 41°C heat threat at 8 mins | 25 |
| | CDN>Krz-RNAi, 12 min | $w^{1118}$, Dcr-2/Y; NP2719-Gal4, RepoGal80/+; UAS-Krz-RNAi/+ | 41°C heat threat at 12 mins | 25 |
| | CDN>Krz-RNAi, 16 min | $w^{1118}$, Dcr-2/Y; NP2719-Gal4, RepoGal80/+; UAS-Krz-RNAi/+ | 41°C heat threat at 16 mins | 24 |
| B | CDN>ChR2-XXM, green-camuiα +Krz-RNAi | $w^{1118}$; NP2719-Gal4, RepoGal80/ChR2-XXM; UAS-green-camuiα/UAS-Krz-RNAi | Not satiated, 10 µM DA | 10 |
| | CDN>ChR2-XXM, green-camuiα +Krz-RNAi-1 | $w^{1118}$; NP2719-Gal4, RepoGal80/ChR2-XXM; UAS-green-camuiα/UAS-Krz-RNAi | Satiated 4.5 hrs, 10 µM DA | 10 |
| C | CDN>Chr + Krz-RNAi Not satiated | $w^{1118}$, Dcr-2/Y; NP2719-Gal4, RepoGal80/+; UAS-Krz-RNAi/UAS-Chrimson | Not satiated, 1 min of 15 µW/mm$^2$ green light at 8 min | 28 |
| | CDN>Chr + Krz-RNAi Satiated | $w^{1118}$, Dcr-2/Y; NP2719-Gal4, RepoGal80/+; UAS-Krz-RNAi/UAS-Chrimson | Satiated 2.5 hrs, 1 min of 15 µW/mm$^2$ green light at 8 min | 26 |
